## Supplementary Data for "Cryo-EM Structures of *Saccharolobus solfataricus* Initiation Complexes with Leaderless mRNAs Highlight Archaeal Features and Eukaryotic Proximity"

**Supplementary Methods**

**Supplementary Tables**

**Supplementary Figures**

**Supplementary References**

**Supplementary Methods**

**Grid inhomogeneous solvation theory analysis of 16S rRNA G889**

Water density around G889 nucleobase of 16S rRNA was studied by explicit water molecular dynamics (MD) simulation, followed by grid inhomogeneous solvation theory (GIST)[1, 2] analysis. GIST is a method giving access to the structure and thermodynamics of solvent in the vicinity of a solute molecule. The ribosomal molecular system was truncated at 12 Å around G889 nucleobase, resulting in 36 RNA nucleotides, 36 experimentally placed water oxygens, and 3 magnesium ions. Two alternative conformations were considered for the 5'-terminal triphosphate nucleotide (A3P) of mRNA, close in space to G889. Hydrogen atoms were added and their positions optimized with the Reduce program[3]. The system was immersed in a truncated octahedron box extending 10 Å away from the solute and filled with water molecules. Supplementary Figure 16A shows the simulated system. Minimization was conducted, starting with the hydrogen atoms alone, then adding the solvent, while gradually reducing the harmonic positional restraints on the rest, until reaching a force constant of 10 kcal/mol/Å$^2$. The reference for positional restraints was the initial conformation. Only heavy atoms of the solute and magnesium ions were restrained. Minimizations were stopped when the RMSD of the gradient components was less than 0.1 kcal/mol/Å. MD simulation was then conducted. Heating to 300 K was first performed, in the constant volume and temperature (NVT) ensemble. Equilibration was then performed in the constant pressure and temperature (NPT) ensemble during 2 ns, then in the NVT ensemble during 10 ns. The production simulation was then run in the NVT ensemble during 100 ns and conformations were saved every 4 ps. The harmonic positional restraints used in final minimization were maintained during MD simulation. The integration time step was 2 fs. Bonds involving hydrogen atoms were constrained with the SHAKE algorithm[4]. Periodic boundary conditions were applied. The Particle Mesh Ewald (PME) method[5] was used to handle long-range electrostatic interactions, while long-range van der Waals interactions were estimated by a continuum model. A cutoff distance of 9 Å was used for direct calculation of interactions. Temperature was controlled with a Langevin thermostat[6], targeting a temperature of 300 K and using a collision frequency of 2 $ps^{-1}$. Pressure was controlled with the Berendsen barostat[7], targeting a pressure of 1 bar and using a relaxation time of 2 ps. The AMBER suite[8] of programs was employed, with the SANDER program for minimization and heating, and the GPU accelerated version of the PMEMD program[9, 10] for equilibration and production. The ff99OL3 force field[11, 12] was used for RNA, along with modrna08[13] for modified nucleotides, Carlson ATP parameters[14] were adapted for the A3P nucleotide, TIP3P[15] was used for water, and Li and Merz parameters[16] (12-6 normal usage set) for magnesium ions. The production simulation trajectory was then analyzed with GIST around G889 nucleobase, as implemented in the CPPTRAJ program[17]. The grid center was defined as the origin of the reference Cartesian frame of G889 nucleobase, following[18]. The grid dimensions were 30, 30, and 36, in the x, y, and z directions (MD simulation axes), while the grid spacing was 0.5 Å. Reference density for water was 0.0329 molecules/Å3. Supplementary Figure 16B shows the simulated system without water molecules, along with water oxygen and hydrogen densities obtained by GIST. Supplementary Figure 16C is a zoomed view on a region above G889 nucleobase. This region is defined along the x', y', z' axes of the reference Cartesian frame [18] of G889 nucleobase, as x'=[-2.5,4.5], y'=[-2.5,2.5], z'=[-4.5,0] Å (gray edges in figure 3). Water oxygen and hydrogen densities are shown to be important (more than 2 times the bulk water density) in the z'=[-4,-2.5] Å region (black edges in Supplementary Figure 16C). This region contains five experimentally placed water molecules (red spheres in Supplementary Figure 16C and Figure 7D). Table 1 presents a selection of water properties calculated with GIST, averaged over these different regions. Properties are water oxygen and hydrogen densities, as well as solute-water and water-water interaction energy densities. These quantitative results show that when gradually focusing on the region above G889 nucleobase, water density increases, as well as both solute-water and water-water interaction energy densities. At the location of the five water molecules that have been placed based on the experimental electronic density, 2 water oxygen and hydrogen densities are 3.75 and 2.00 times higher than bulk water density, while solute-water and water-water interaction energy densities are -1.58 and -0.60 kcal/mol/Å3. Robustness and convergence of the GIST analysis was assessed by redoing the MD simulations without the 36 experimentally placed water oxygens. The results obtained are almost identical to those obtained including experimental water oxygens, indicating that initial placement of water molecules has limited influence and suggesting that the MD simulation has sufficiently sampled the water conformational space. Overall, these results support the assignment of the five water molecules in the region above G889 nucleobase. Favorable interaction energies of these water molecules with G889 is calculated. These water positions are compatible with OH and/or lone pair stacking interactions with G889, as studied by[18] with quantum chemical calculations. At last, we note that the shape of water density above G889 nucleobase is suggestive of a purine nucleobase, where the cycle atoms would be replaced by a network of water molecules.

### Supplementary Tables

| mRNAs | Size | Sequences 5'>3' | Production-purification |
|---|---|---|---|
| model-SD | 658 | GGGAGACCACAACGGUUUCCCTCT AGAAAGUAGGGUUAGGCAGAAAUU UGGAGGUGAUUUAAAUGCCAAAGG AGAAGCCCCACGUUAACAUCGUGU UUAUCGGACACGUAGACCACGGAA AGAGCACGACCAUCGGAAGGCUCC UCUACGACACCGGGAACAUCCCAG AGACCAUCAUCAAGAAGUUCGA… | [19] |
| Ss-aEF1A-like | 182 | GUUAAAGUGAUUUGAUGUAAUGA AAUUACCCCGAGAGAAUGAUGAAG GAAAAGUAGAAUAUAAGCUUAUUC UUUCAAGUGUAACCCCGGAUCGUC UUCAAGAAUUAGCUACUCAAAUGA AGUAUAGGUUAGAGGAAGGUGAU GGUGAAGCAUUUUACGUUAUAGGU GUAAGUGAUGAAGGCG | Cloned into pET3a BglII-XhoI<br>In vitro transcription<br>MonoQ |
| Ss-MAP | 161 | 5'-OH- AUGACUGAGGAUGAACUCAAUAAG CUUUUAUUAGCAGGUAAGAUUGCA GCUAAGGCUAGAGAUGAAGUUUCA UUAGACGUUAAAGCUAGUGCUAAG GUUUUAGAUAUUUGUGAAGAGGU UGAAAGUAUAAUAAUCGAAAAUAA AGCGUUUCCAUCAUUUCC | Cloned into pET3a XbaI-XhoI<br>5' hammerhead ribozyme<br>In vitro transcription<br>MonoQ |
| Ss-aIF2β | 171 | 5'PPP- GUGAGUUCAGAAAAAGAAUACGUA GAAAUGCUUGAUAGGCUAUACUCG AAAUUACCAGAAAAAGGACGUAAG GAAGGUACACAAUCAUUGCCUAAU AUGAUAAUACUCAAUAUAGGAAAU ACUACUAUAAUUAGAAACUUUGCG GAGUAUUGUGAUAGAAUUAGAAG AGAGG | Cloned into pET3a BglII-XhoI<br>In vitro transcription<br>MonoQ |
| model-SD (short) | 28 | 5'-OH- AUUUGGAGGUGAUUUAAAUGCCAA AG | Dharmacon |
| Ss-aEF1A-like (short) | 24 | 5'-OH- UAAAGUGAUUUGAUGUAAUGAAAU UACC | Eurogentec |
| Ss-MAP (short) | 15 | 5'-PPP-AUGACUGAGGAUGAA | Eurogentec |
| Ss-aIF2β (short) | 15 | 5'-PPP-GUGAGUUCAGAAAAA | In vitro transcription. MonoQ |

**Supplementary Table 1: model mRNAs used in the toeprinting experiments and cryo-EM studies**
The Shine-Dalgarno sequence complementary to the anti-SD sequence is colored in red when present. The start codon and the base corresponding to the major toeprinting signal is colored in blue in long mRNAs. Pa-aEF1A, Ss-MAP, Ss-aEF1A-like mRNAs were chemically synthetized (Eurogentec). A 5' triphosphate group was chemically added to Ss-aEF1A-like mRNA (Eurogentec). Ss-aIF2β mRNA was in vitro transcribed (see Methods). The violet bases in Ss-MAP short lmRNA indicate the pseudo-SD sequence located downstream from the start codon.

| | DS1 | DS1 | DS2 | DS2 | DS2 | DS3 | DS3 | DS4 |
|---|---|---|---|---|---|---|---|---|
| **Data collection** | IC2-down | IC2-up | 30S-HR | IC2-down | IC2-up | IC2-down | IC2-up | IC2-down |
| PDB ID | 9FRK | 9FRL | 9FHL | 9FRA | 9FSF | 9FY0 | 9FS6 | 9FS8 |
| EMDB ID | 50716 | 50717 | 50445 | 50709 | 50727 | 50854 | 50724 | 50725 |
| Microscope | ESRF | ESRF | TFS Krios Pasteur | TFS Krios Pasteur | TFS Krios Pasteur | TFS Krios Pasteur | TFS Krios Pasteur | Glacios 2 Pasteur |
| Camera | Gatan K3 | | Gatan K3 | | | Falcon4i | | Falcon4i |
| Magnification | 130000 | | 105000 | | | 165000 | | 130000 |
| Voltage (kV) | 300 | | 300 | | | 300 | | 200 |
| Electron exposure (e–/Å$^2$) | 40 | | 40 | | | 20 | | 40 |
| Defocus range (μm) | -2.9 to -0.9 | | -3 to -1 | | | -2.6 to -0.8 | | -2.6 to -0.8 |
| Pixel size (Å) | 1.053 | | 0.86 | | | 0.73 | | 0.88 |
| Initial particle (no.) | 269 000 | 269 000 | 975 000 | 975 000 | 975 000 | 234 000 | 234 000 | 308 783 |
| Final particle (no.) | 28 000 | 38 000 | 766 000 | 62 000 | 108 000 | 43 000 | 49 000 | 156 |
| Resolution (unmasked. Å) | 4.03 | 3.73 | 2.87 | 3.01 | 3.33 | 4.09 | 4 | 4.2 |
| Resolution (masked. Å) | 3.02 | 3 | 2.52 | 2.86 | 2.85 | 2.92 | 2.92 | 3.72 |
| FSC threshold | 0.143 | 0.143 | 0.143 | 0.143 | 0.143 | 0.143 | 0.143 | 0.143 |
| **Refinement** | | | | | | | | |
| d FSC model (Å). threshold 0.5 | 3.1 | 3.4 | 2.6 | 3 | 2.9 | 2.9 | 2.9 | 3.9 |
| Map sharpening B factor (Å$^2$) | - | - | -96.41 | -10 | -10 | - | - | - |
| Model composition | | | | | | | | |
| Non-hydrogen atoms | 66394 | 65224 | 64047 | 66676 | 65783 | 66885 | 65634 | 66643 |
| Protein residues | 3937 | 3937 | 393 | 3938 | 3936 | 4026 | 4029 | 4028 |
| Nucleotides | 1593 | 1540 | 1460 | 1597 | 1552 | 1574 | 1520 | 1579 |
| Ions and spermines | ZN:8 MG:59 SPM:36 | ZN:8 MG:53 SPM:31 | ZN:8 MG:59 SPM:44 | ZN:8 MG:58 SPM:42 | ZN:8 MG:64 SPM:44 | ZN:9 MG:47 SPM:43 | ZN:9 MG:57 SPM:39 | ZN:9 MG:57 SPM:30 |
| Water molecules | 165 | 210 | 515 | 221 | 276 | 273 | 199 | 91 |
| Average B factors (Å$^2$) | | | | | | | | |
| Protein | 104.9 | 80.5 | 31.8 | 98.3 | 94.8 | 83.2 | 86.8 | 148 |
| Nucleic acid | 106.8 | 106.8 | 30.8 | 112.8 | 101.9 | 77.8 | 75.9 | 149.7 |
| Ligand | 83.2 | 81.8 | 25.1 | 85.3 | 84.4 | 67.5 | 64.4 | 127 |
| Water molecules | 73.5 | 77.1 | 20.3 | 78.2 | 79.22 | 53.7 | 50.1 | 122.8 |
| R.m.s. deviations | | | | | | | | |
| Bond lengths (Å) | 0.009 | 0.005 | 0.007 | 0.004 | 0.005 | 0.005 | 0.008 | 0.008 |
| Bond angles (°) | 0.983 | 0.813 | 0.918 | 0.627 | 0.64 | 0.683 | 0.857 | 0.925 |
| **Validation** | | | | | | | | |
| MolProbity score | 2.3 | 2.3 | 2.3 | 2.2 | 2.2 | 2.2 | 2.2 | 2.28 |
| Clashscore | 19.38 | 18.17 | 13.7 | 12.4 | 13 | 16.2 | 16 | 23.87 |
| Poor rotamers (%) | 2 | 2.38 | 3.3 | 2.5 | 2.5 | 2 | 2.1 | 1.52 |
| Ramachandran plot | | | | | | | | |
| Favored (%) | 95.74 | 95.72 | 95.92 | 96.23 | 95.97 | 95.78 | 95.76 | 96.06 |
| Allowed (%) | 4.23 | 4.26 | 4.02 | 3.74 | 4 | 4.17 | 4.21 | 3.94 |
| Disallowed (%) | 0.03 | 0.03 | 0.05 | 0 | 0.03 | 0.05 | 0.03 | 0 |
| Correlation coefficients | | | | | | | | |
| Mask CC | 0.87 | 0.8 | 0.84 | 0.9 | 0.9 | 0.86 | 0.86 | 0.85 |
| Volume CC | 0.87 | 0.8 | 0.81 | 0.9 | 0.9 | 0.85 | 0.82 | 0.84 |

**Supplementary Table 2: Cryo-EM data collection, refinement and validation statistics**

| RNA modification | | Position(s) in *S. solfataricus* 16S rRNA | Position(s) in *P. abyssi* 16S rRNA | Position(s) in *T. kodakarensis* 16S rRNA |
|---|---|---|---|---|
| $ac^4C$ | N4-acetylcytidine | 1466,1467, 1477*,1478* | 17,53,286,303, 319,379,394, 479,511,546, 590,626,636, 703,718,731, 751,828,839, 848,851,868, 957,1028, 1147,1233, 1239,1479 | 5,41,136,298,373 ,458,501,525, 569,605,697,730, 818,827,847, 1007,1217,1459 |
| Am | 2'-O-methyladenosine | 494* | 373 | 352 |
| Cm | 2'-O-methylcytidine | 246,313,481,1060, 1366 | 129,846,1036,1040, 1376 | 190,296,535,752,825, 1354,1361 |
| Gm | 2'-O-methylguanosine | 337,399,672,927*, 1018*,1061,1194 | 467,471,519,657, 680,873,913,934, 1069 | 138,246,320,381,346,350, 446,450,498,532,636,659, 771,852,876,892,913,994, 1025,1026,1107,1265,1275, 1434 |
| $m^5C$ | 5-methylcytidine | 1368 | 535,693,875,1025, 1202,1374,1496, 1498,1505 | 457,464,672,854,942, 1004,1276,1374,1476, 1478 |
| $m^6A$ | N6-methyladenosine | 1457* | 1469 | 1449 |
| $m^{6,6}A$ | N6.N6-dimethyladenosine | 1475* | 1487,1488 | 1467,1468 |
| Um | 2'-O-methyluridine | 52,875*,1032,1344* | 20,64,774,787,830, 1177,1380 | 8,52,240,304,479,1358 |
| $m^1acp^3\psi$ | 1-methyl-3-α-amino-α-carboxyl-propyl pseudouridine | 930* | | |

**Supplementary Table 3** : **Modified residues localized in 16S rRNA sequence of *S. solfataricus* and compared to *P. abyssi* and *T. kodakarensis***
The name of the rRNA modification and its position in 16S rRNA sequences are indicated. The modifications colored in red are observed in the three archaea. The asterisk indicates that the presence of a post-transcriptional modification at this position was observed in[20]. Reference structures are those of *P. abyssi* 16S rRNA (7ZHG)[21] and of *T. kodakarensis* (6TH6)[22].

| Compounds | Elemental composition ($MH^+$) | $MH^+$ (*m/z*) | $BH^+$ (*m/z*) | $2MH^+$ (*m/z*) | tr (min) |
|---|---|---|---|---|---|
| C | $C_9H_{14}N_3O_5$ | 244.0928 | 112.050 | 487.17 | 1.3 |
| $m^5C$ | $C_{10}H_{16}N_3O_5$ | 258.1084 | 126.066 | - | 1.6 |
| Cm | $C_{10}H_{16}N_3O_5$ | 258.1084 | 112.050 | 515.2091 | 2.2 |
| U | $C_9H_{13}N_2O_6$ | 245.0768 | 113.035 | 489.1460 | 2.4 |
| A | $C_{10}H_{14}N_5O_4$ | 268.1040 | 136.061 | - | 3.0 |
| G | $C_{10}H_{14}N_5O_5$ | 284.0989 | 152.056 | 567.1901 | 3.6 |
| Am | $C_{11}H_{16}N_5O_4$ | 282.1196 | 136.061 | - | 4.3 |
| Gm | $C_{11}H_{16}N_5O_5$ | 298.1146 | 152.056 | 595.2227 | 4.8 |
| mG | $C_{11}H_{16}N_5O_5$ | 298.1146 | 166.07 | | *4.9* |
| $m^6A$ | $C_{11}H_{16}N_5O_4$ | 282.1196 | 150.077 | - | 4.9 |
| $ac^4C$ | $C_{11}H_{16}N_3O_6$ | 286.1033 | 154.061 | - | 5.2 |
| $m^{6,6}A$ | $C_{12}H_{18}N_5O_4$ | 296.1353 | 164.093 | - | 7 |

**Supplementary Table 4: Detection of nucleosides in *S. solfataricus* 16S rRNA.**
For each detected nucleoside. the retention time is indicated (See Methods). The measured molecular mass is indicated for the $MH^+$ nucleoside and for the $BH^+$ ion (nucleoside fragment ion derived from the base moiety). *Low intensity; nd : not detected. Gm was detected but not localized in the cryo-EM maps. The presence of multimers (2MH+) reveals higher concentration. only the $MH^+$ nucleoside for $m_1acp^3\psi$ was detected

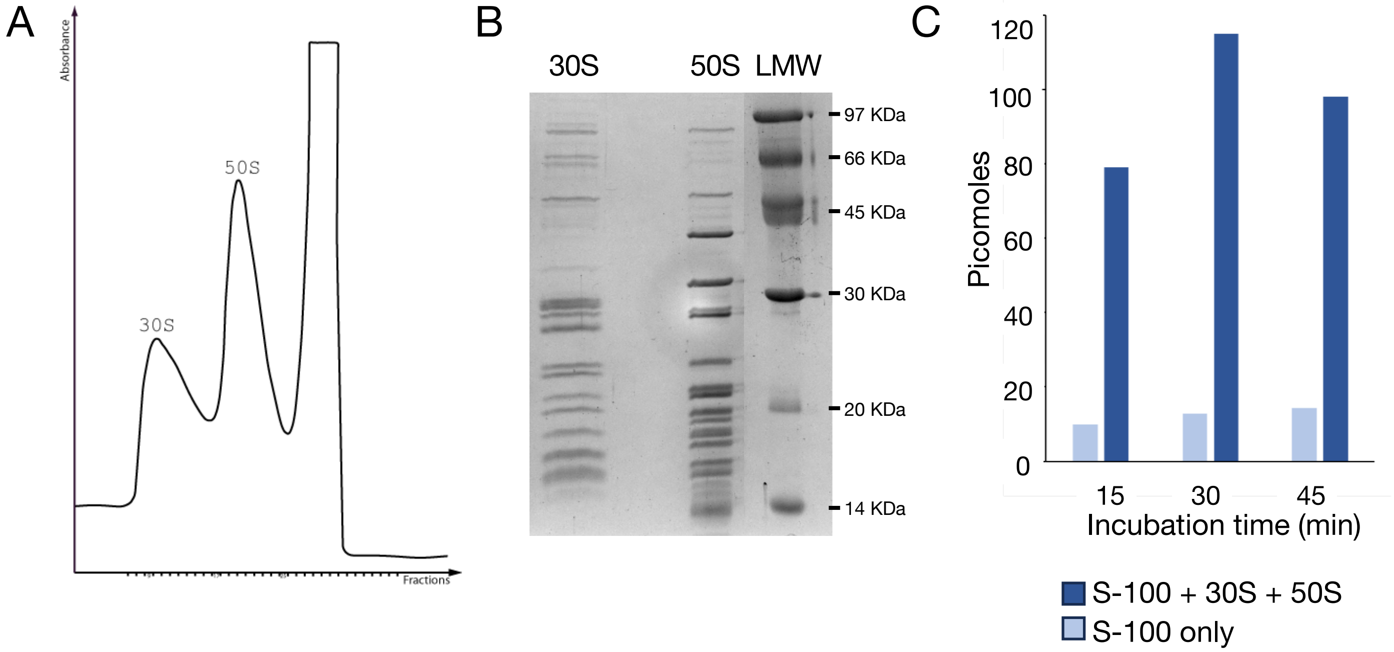

**Supplementary Figure 1: Preparation of *S. solfataricus* ribosomal subunits**

A. Sucrose gradient (10-30%) profile of the ribosomal subunits in buffer A.
B. SDS-PAGE analysis 30S and 50S preparations. Lane 1: 30S, lane 2: 50S. This 30S preparation was analyzed by mass spectrometry.
C. In vitro translation assay. In vitro poly-U translation conditions were adapted from[23,24], see Methods. The graph shows the amount of picomoles of [$^3$H]Phe polymerized after the indicated times of incubation, in the presence of S-100 fraction only (light blue bars) or in the presence of S-100 plus purified 30S and 50S (dark blue bars). The experiment shows that the purified ribosomal subunits are active in poly-Phe synthesis, being able to polymerize 1 pmol phe/pmol 50S particles at 60°C, in reasonable agreement with[24].

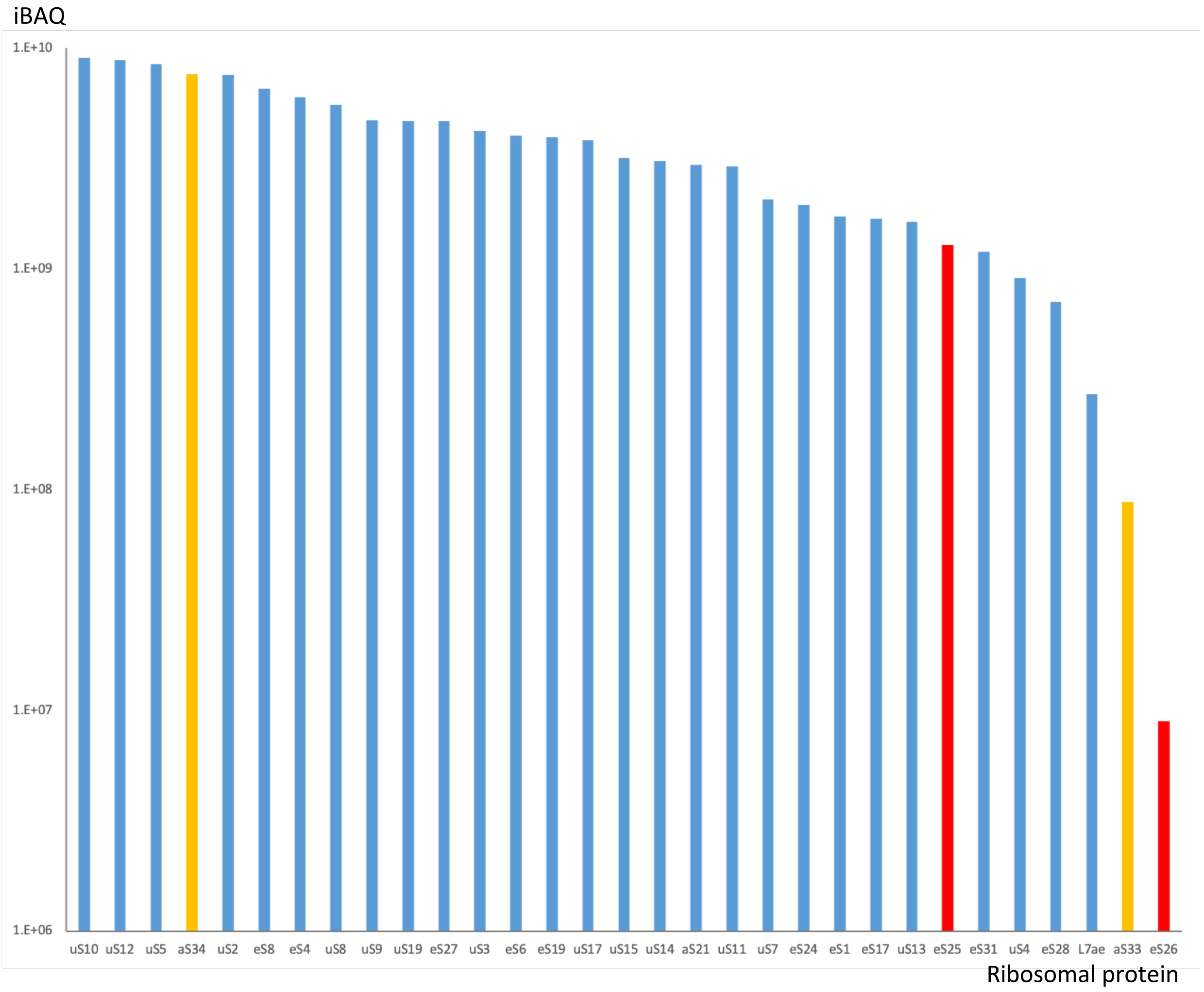

**Supplementary Figure 2: Ribosomal proteins identified by nanoLC-MS/MS analysis**

Mass spectrometry analysis (see methods) was performed on the 30S sample shown in Supplementary Figure 1B. Data processing was performed with MaxQuant software against a database containing the *S. solfataricus* P2/DSM1617 proteome (Uniprot), the eS26 sequence (tr|A0A0E3MCW1|A0A0E3MCW1_SACSO;Q97ZR1) and that of D0KTI0_SACS9 (named here aS33) which were not in the *S. solfataricus* library. iBAQ intensities are plotted in a log scale for each ribosomal protein. aS33 and aS34 are shown with orange bars. eS25 and eS26 are shown with red bars. eS30, observed in our 30S cryo-EM structures, was not detected in this nanoLC-MS/MS analysis likely because of its small size and high-content of lysine and arginine residues (9 K, 7 R out of 51 residues, leading only to three potentially detectable peptides).

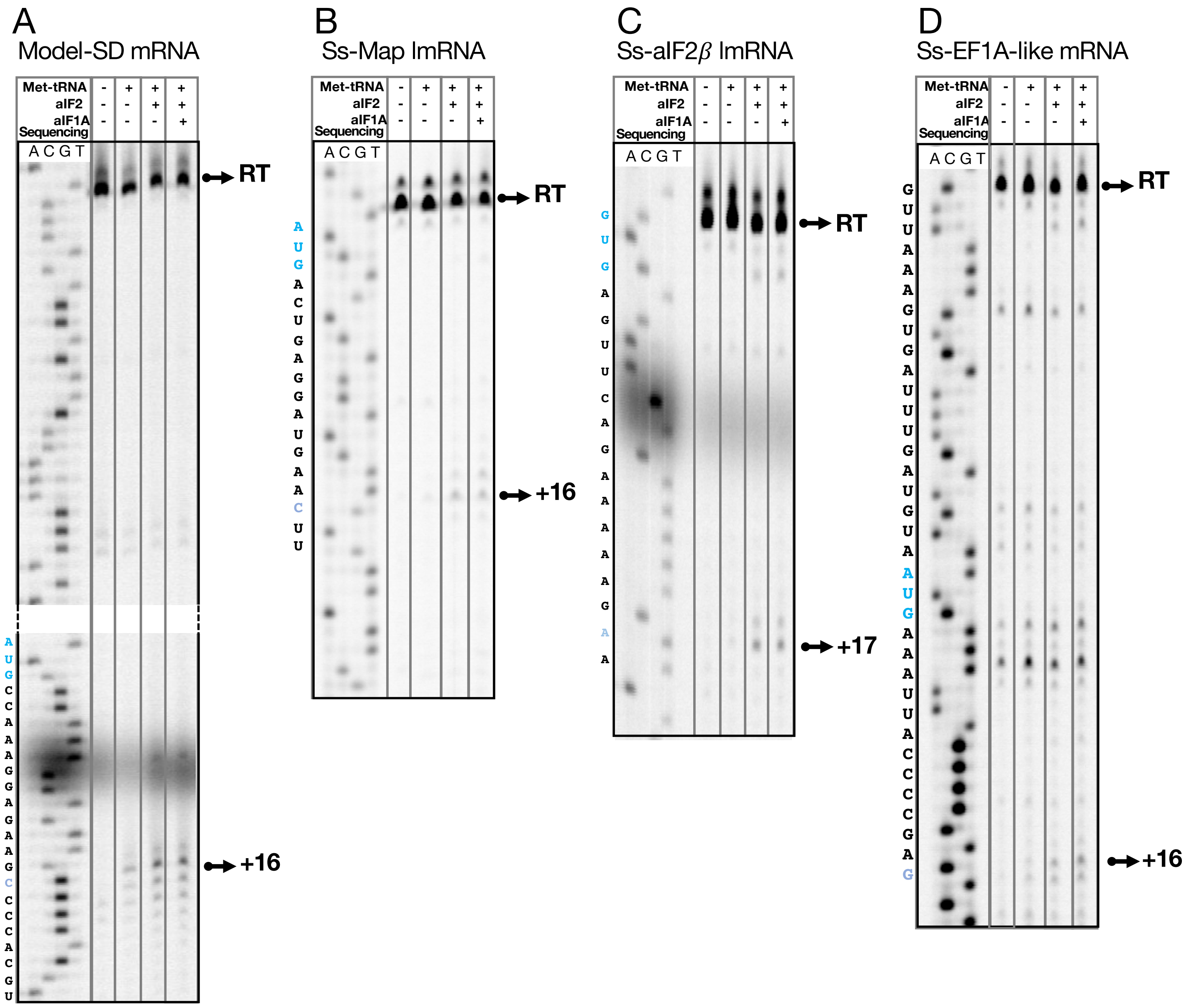

**Supplementary Figure 3: Examples of toeprinting analysis of 30S complexes assembled on various mRNAs in the presence of the indicated initiation factor and of Met-tRNA$_i^{Met}$**

A. Typical toeprinting experiment using model-SD mRNA. The mRNA sequence complementary to the DNA sequence is shown on the left of the gel image. The positions of the start codon and of the main toeprinting signals are indicated in blue. Full-length cDNA is marked RT, Lane 1: toeprinting signal for a 30S:mRNA complex, lane 2: 30S:mRNA:Met-tRNA$_i^{Met}$, lane 3: 30S:mRNA-Met-tRNA$_i^{Met}$:aIF2, lane 4: 30S:mRNA-Met-tRNA$_i^{Met}$:aIF2:aIF1A (see also Figure 1A).
B. Same as panel A but using the Ss-Map lmRNA.
C. Same as panel A but using the Ss-aIF2β lmRNA.
D. Same as panel A but using the Ss-EF1A-like mRNA.

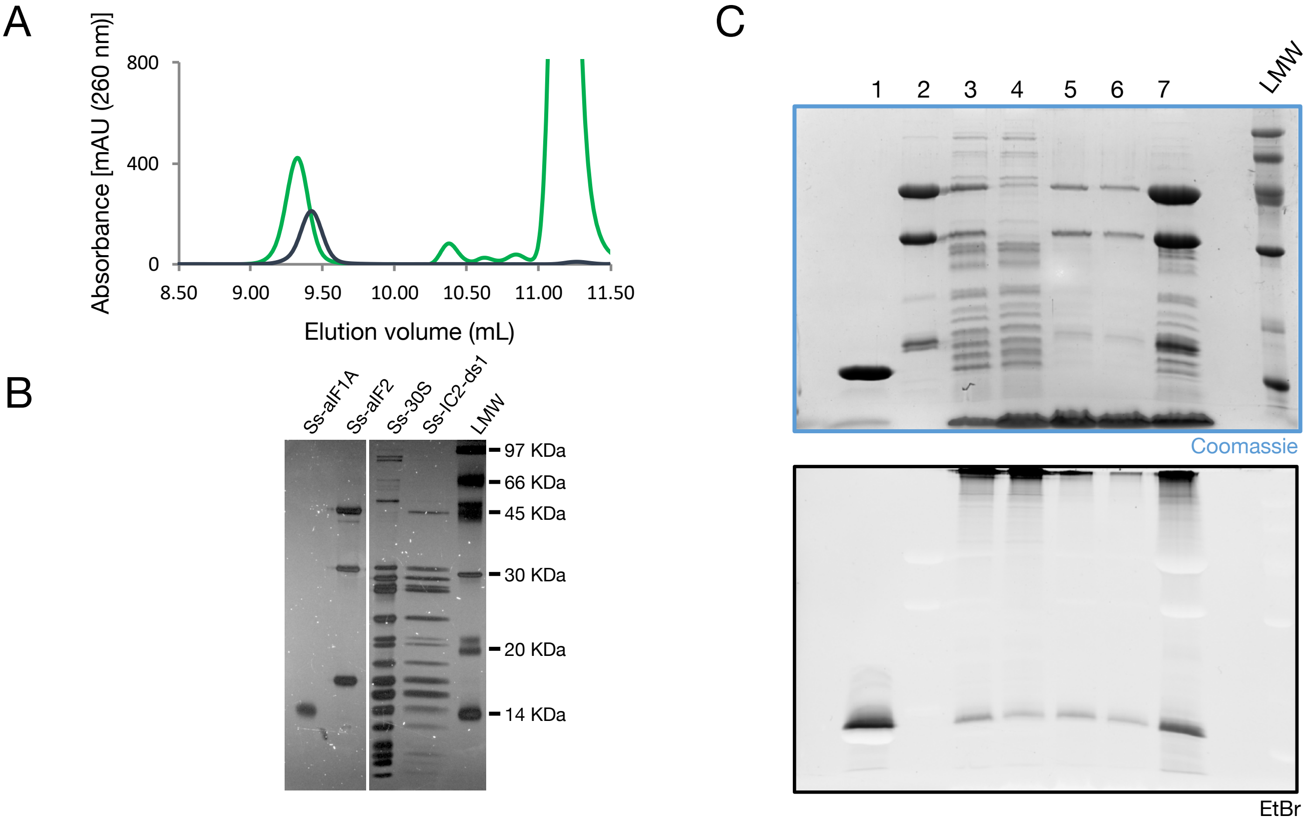

**Supplementary Figure 4: Preparation of initiation complexes**

A. Bio-Agilent Sec5 chromatograms for Ss-30S (black) and model-SD mRNA initiation complex (IC) used to collect dataset 1 (green).
B. SDS-PAGE analysis of Ss-aIF1A, Ss-aIF2, Ss-30S, and Ss-IC2-DS1 purified by size exclusion chromatography. The gel was stained using SYPRO rubis (Thermo Fisher).
C. SDS-PAGE analysis of affinity purification steps of IC used to collect dataset 2. The upper part of the gel was stained with Coomassie blue (up) and ethidium bromide (bottom) to reveal the initiator tRNA. Lane 1: mixture of Ss-aIF1A and Met-tRNA, lane 2: Ss-aIF2, lane 3: mixture of Ss-30S + Ss-Map-lmRNA + Ss-aIF1A + Ss-aIF2 + Met-tRNA$_i^{Met}$ , lane 4: Flow-through fraction, lanes 5-6: elution fractions, lane 7: concentrated sample.

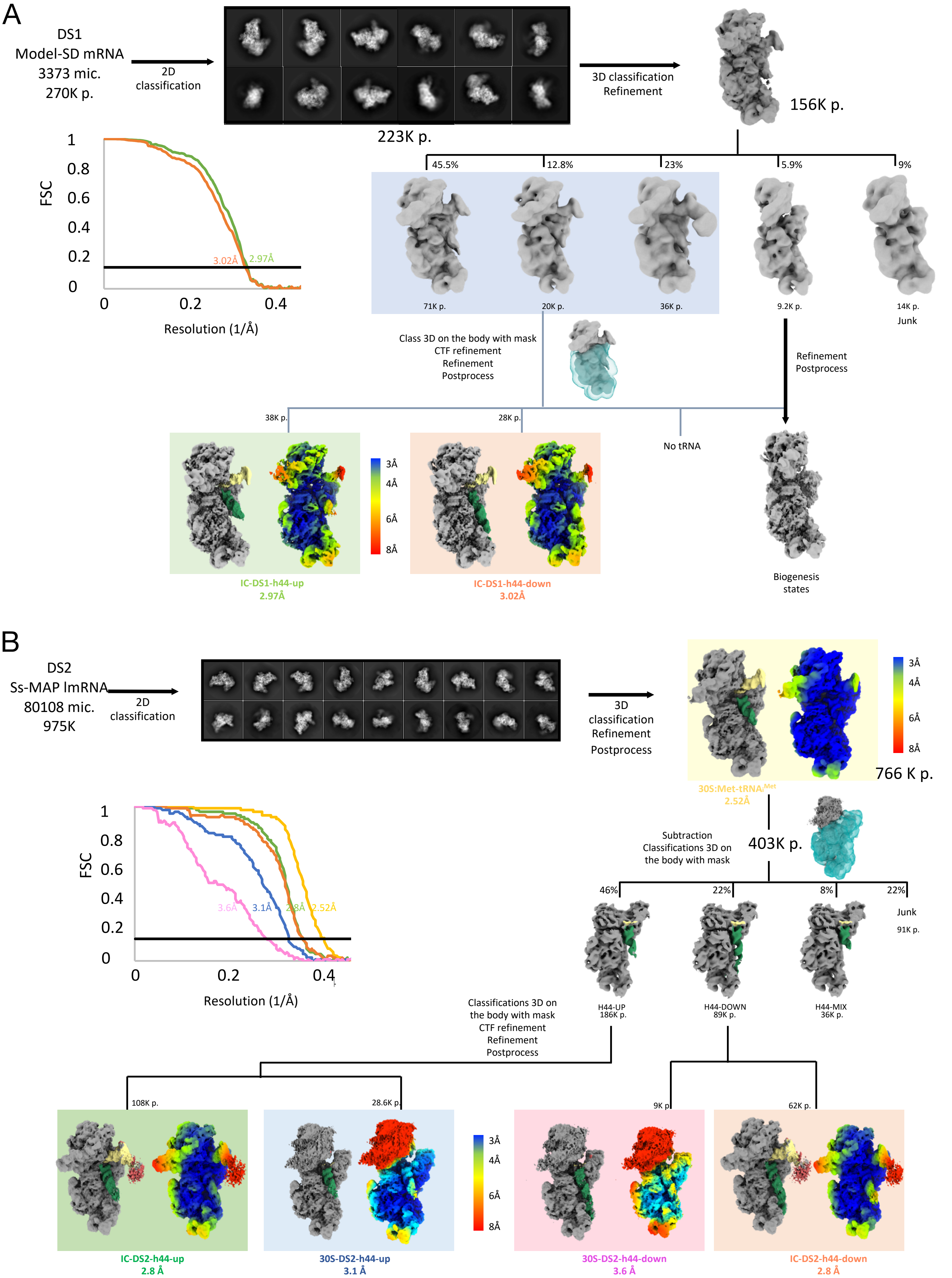
A
DS1
Model-SD mRNA
3373 mic.
270K p.
2D classification
223K p.
3D classification
Refinement
156K p.
FSC
Resolution (1/Å)
3.02Å
2.97Å
45.5%
12.8%
23%
5.9%
9%
71K p.
20K p.
36K p.
9.2K p.
14K p.
Junk
Class 3D on the body with mask
CTF refinement
Refinement
Postprocess
Refinement
Postprocess
38K p.
28K p.
No tRNA
3Å
4Å
6Å
8Å
IC-DS1-h44-up
2.97Å
IC-DS1-h44-down
3.02Å
Biogenesis states
B
DS2
Ss-MAP lmRNA
80108 mic.
975K
2D classification
3D classification
Refinement
Postprocess
766 K p.
30S:Met-tRNAi Met
2.52Å
Subtraction
Classifications 3D on the body with mask
403K p.
46%
22%
8%
22%
Junk
91K p.
H44-UP
186K p.
H44-DOWN
89K p.
H44-MIX
36K p.
3.6Å
3.1Å
2.8Å
2.52Å
Classifications 3D on the body with mask
CTF refinement
Refinement
Postprocess
108K p.
28.6K p.
9K p.
62K p.
IC-DS2-h44-up
2.8 Å
30S-DS2-h44-up
3.1 Å
30S-DS2-h44-down
3.6 Å
IC-DS2-h44-down
2.8 Å

C

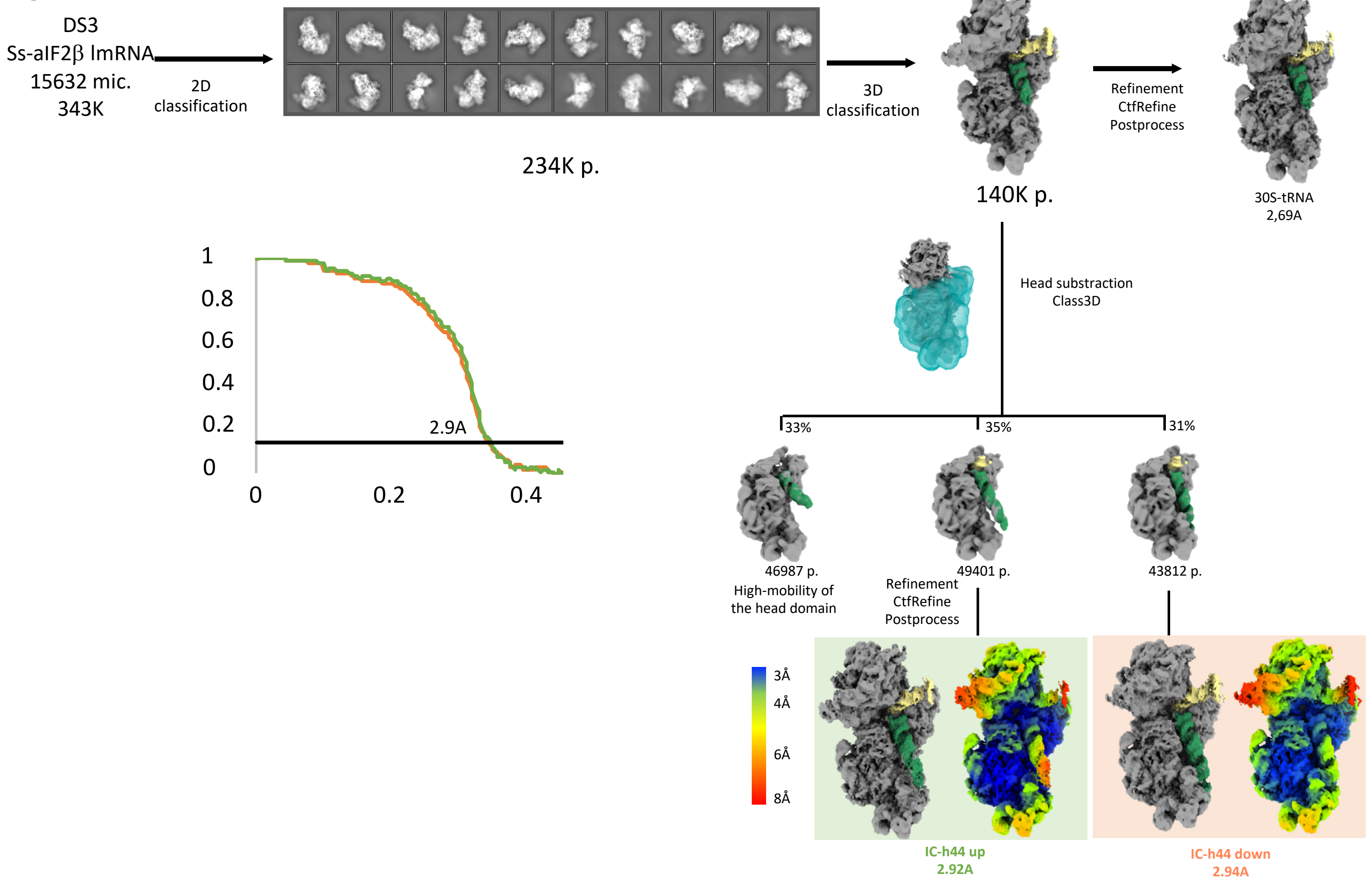
DS3
Ss-aIF2β lmRNA
15632 mic.
343K
2D
classification
234K p.
3D
classification
140K p.
Refinement
CtfRefine
Postprocess
30S-tRNA
2,69A
Head substraction
Class3D
33%
35%
31%
46987 p.
High-mobility of
the head domain
49401 p.
Refinement
CtfRefine
Postprocess
43812 p.
1
0.8
0.6
0.4
0.2
0
0
0.2
0.4
2.9A
3Å
4Å
6Å
8Å
IC-h44 up
2.92A
IC-h44 down
2.94A

D

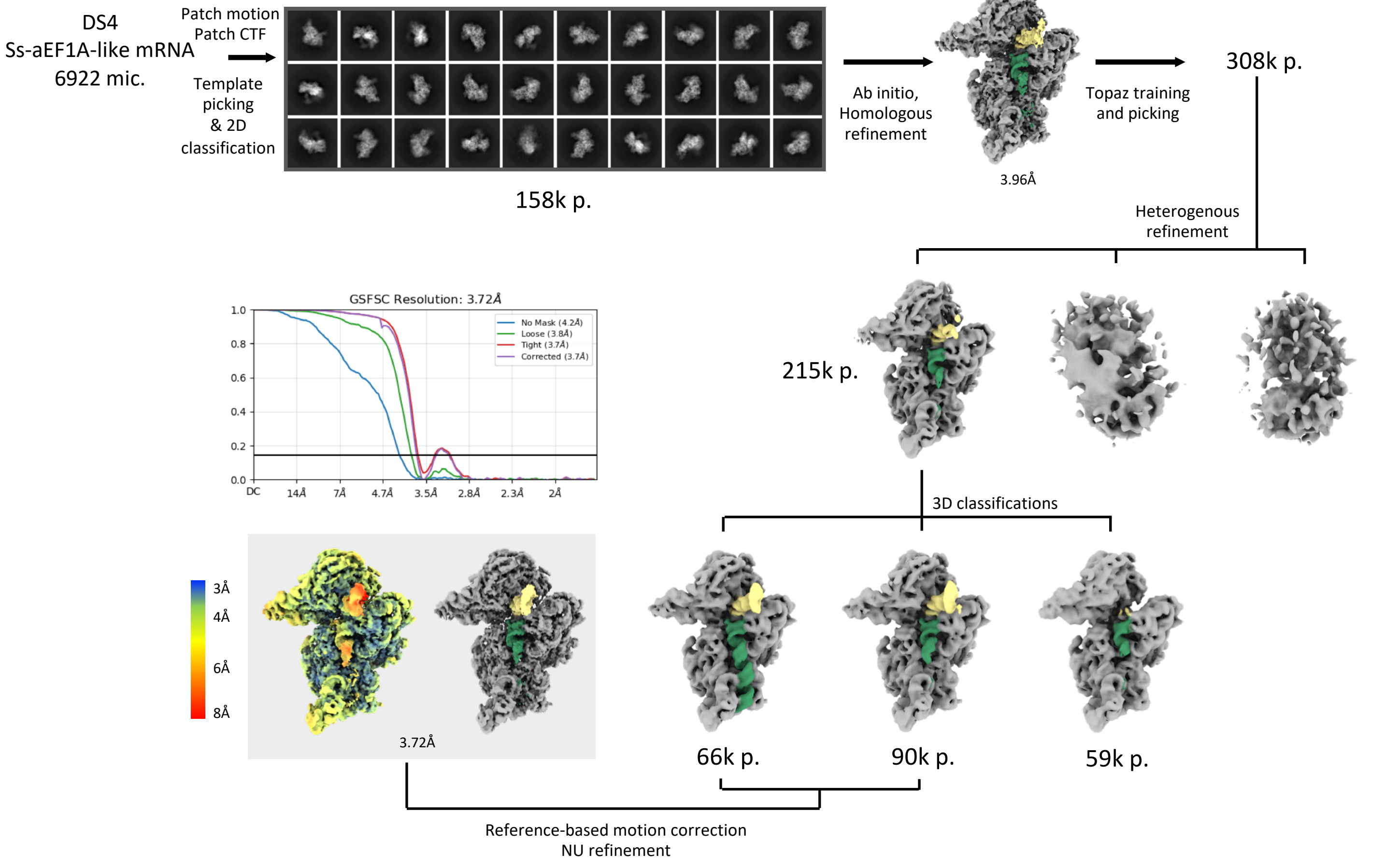
DS4
Ss-aEF1A-like mRNA
6922 mic.
Patch motion
Patch CTF
Template
picking
& 2D
classification
158k p.
Ab initio,
Homologous
refinement
3.96Å
Topaz training
and picking
308k p.
Heterogenous
refinement
215k p.
GSFSC Resolution: 3.72Å
No Mask (4.2Å)
Loose (3.8Å)
Tight (3.7Å)
Corrected (3.7Å)
1.0
0.8
0.6
0.4
0.2
0.0
DC
14Å
7Å
4.7Å
3.5Å
2.8Å
2.3Å
2Å
3D classifications
3Å
4Å
6Å
8Å
3.72Å
66k p.
90k p.
59k p.
Reference-based motion correction
NU refinement

**Supplementary Figure 5: Cryo-EM data processing**

A. Flowcharts of cryo-EM data processing of dataset 1 (DS1) corresponding to an initiation complex assembled on the model-SD mRNA. Data processing was conducted in Relion[25]. Cryo-EM maps filtered and coloured by local resolution are shown. To sort the h44-up and –down conformations, 3D classification was done with a mask containing the 30S body and the possible conformations of h44. This mask is shown as a blue transparent surface. Fourier shell correlation curves obtained by masking the two half maps and calculated the cross-correlation between the masked volumes. Resolution was estimated using the 0.143 cutoff criterion. The green curve is for the IC-DS1-h44-up complex and the orange curve is for the IC-DS1-h44-down complex.
B. Same analysis for dataset 2 (DS2). The IC complex is assembled on Ss-MAP lmRNA. To sort the h44-up and -down conformations, the density of the 30S head was first subtracted. The subtracted particles were then submitted to 3D classification with a mask encompassing the 30S body, the tRNA and the possible conformations of h44. Fourier shell correlation curves are as follows: yellow for 30S:Met-$tRNA_i^{Met}$, green for IC-DS2-h44-up, orange for IC-DS1-h44-down, blue for 30S-DS2-h44-up, violet for 30S-DS2-h44-down.
C. Same analysis for dataset 3 (DS3). The IC complex is assembled on aIF2β-lmRNA. The strategy for sorting the h44-up and -down conformations was the same as for DS2. Fourier shell correlation curves are as follows: green for IC-DS3-h44-up, orange for IC-DS3-h44-down.
D. Data processing of dataset 4 (DS4). The IC complex is assembled on EF1A-like mRNA. Data processing was conducted in cryosparc 4.0[26].

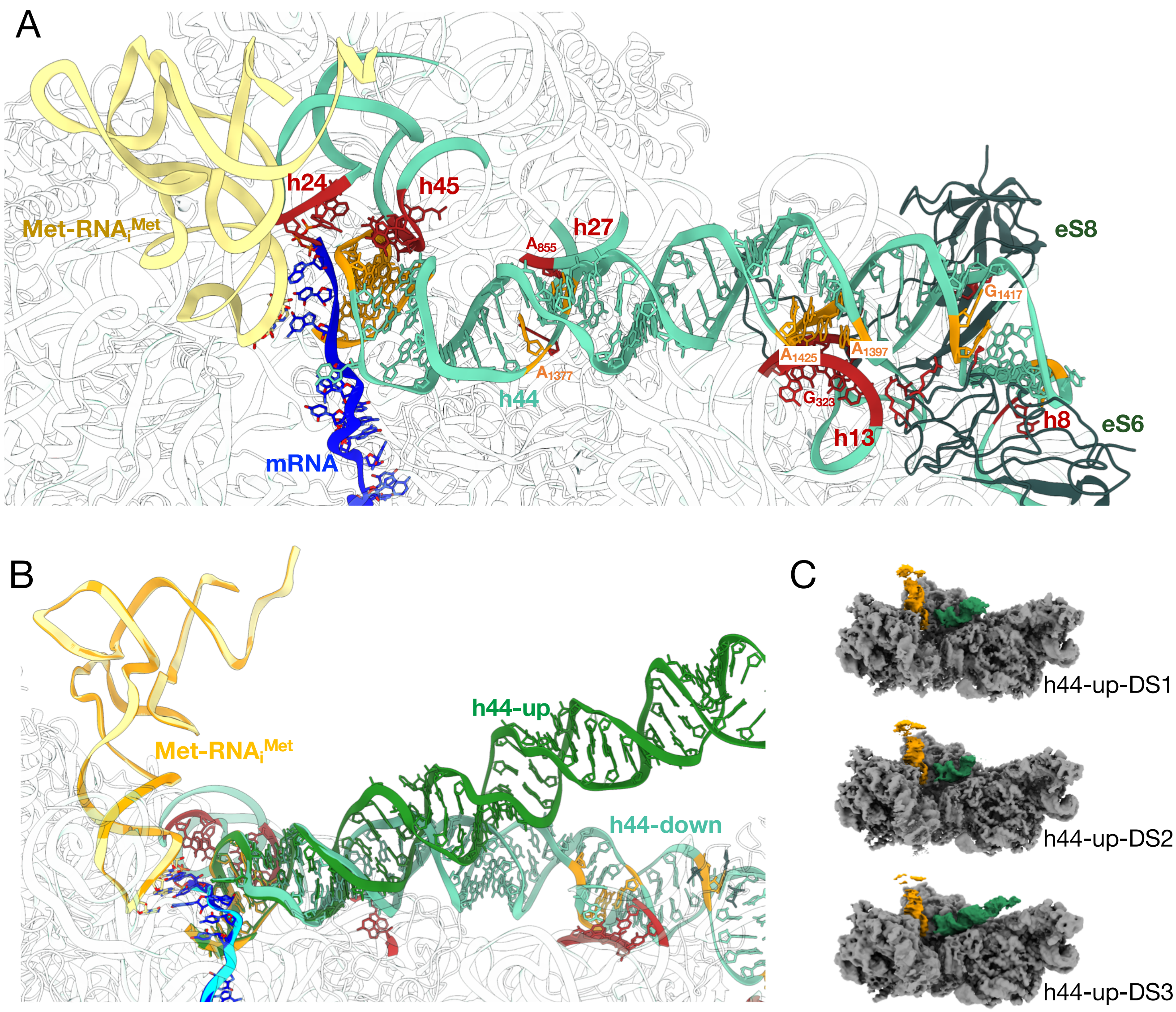

**Supplementary Figure 6: Contacts between h44 and 30S**

A. The upper part of h44 and residues involved in interactions with the body of the 30S are shown in orange. The mRNA is in dark blue and the initiator Met-tRNA is in yellow. The upper part of h44 interacts with h24 and h45 and forms tRNA binding sites. A1377 and C1379 interact with A872 and A855 in h27 bulge; A1397 and A1425 interact with the minor groove of h13 via A-minor interactions with residues G323 and C340. The last bulge of the h44 helix is packed onto h8 minor groove. Two ribosomal proteins interacts with h44, eS8 and eS6.
B. Superimposition of the h44-up and h44-down IC-DS2 cryo-EM structures. The h44-down structure is shown with the same color as in panel A. For the h44-up structure, h44 is in green, the tRNA is in orange and the mRNA is in cyan. The view shows that the conformation of the upper part of h44 is not changed in the up or in the down structures.
C. Cryo-EM maps for DS1, DS2 and DS3 showing h44-up positions (green) and the tRNA.

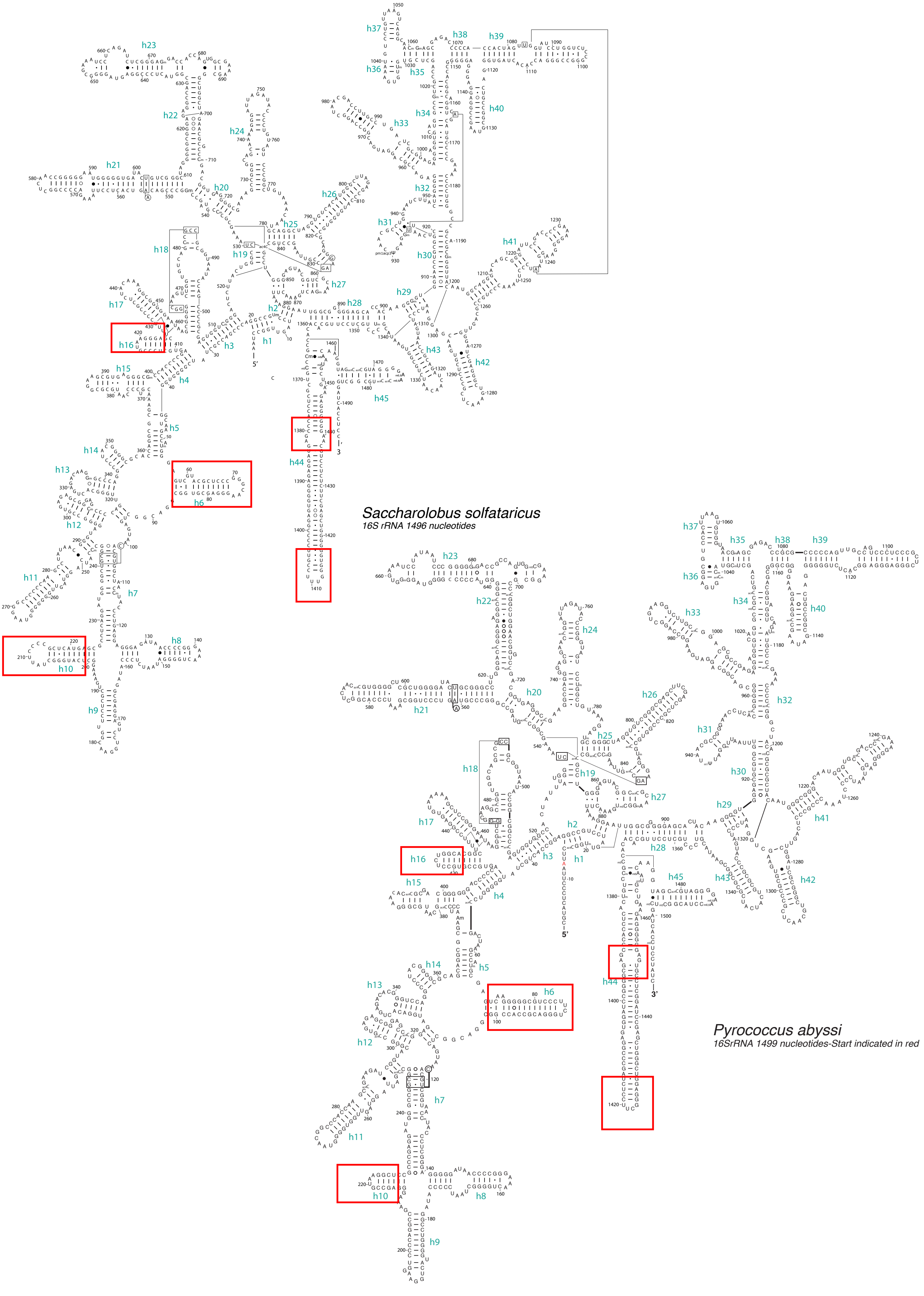

**Supplementary Figure 7: Comparison of 16S rRNAs from S. *solfataricus* and *P. abyssi***

Secondary structure diagrams were retrieved from https://crw-site.chemistry.gatech.edu/ and updated according the cryo-EM structures. The 5’ end of *P. abyssi* 16S rRNA observed in the cryo-EM structure is indicated with a red letter.

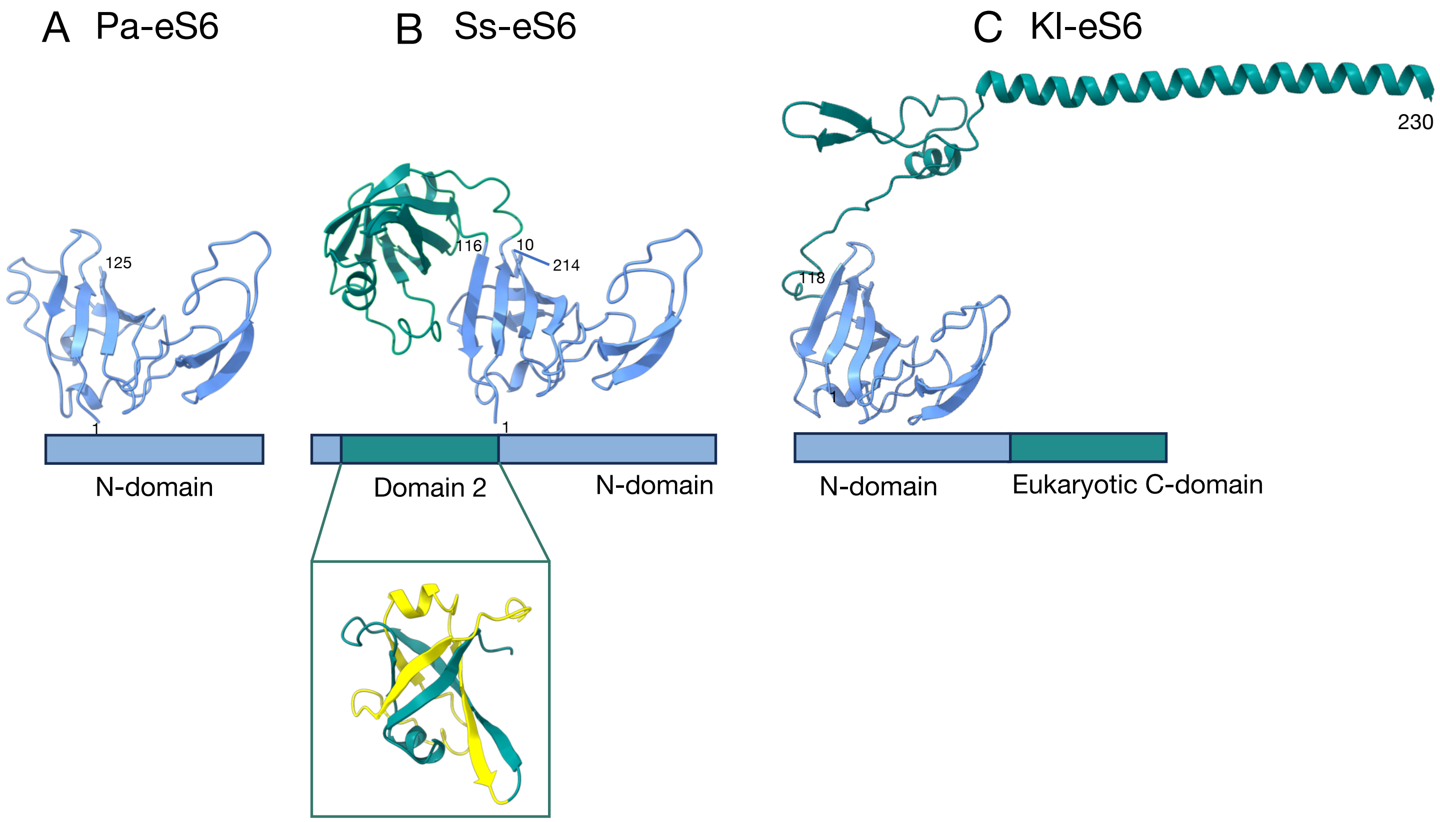

**Supplementary Figure 8: Archaeal and eukaryotic eS6**

A. *P. abyssi* eS6 (PDB 7ZHG)[21].

B. S. *solfataricus* eS6 (present study). A cartoon of *S. solfataricus* eS6 domain 2 showing the double-γ-β-barrel made up of two pseudo-symmetric ββαβ units is also shown.

*C.* *K. lactis* eS6 (PDB 6FYX)[27].

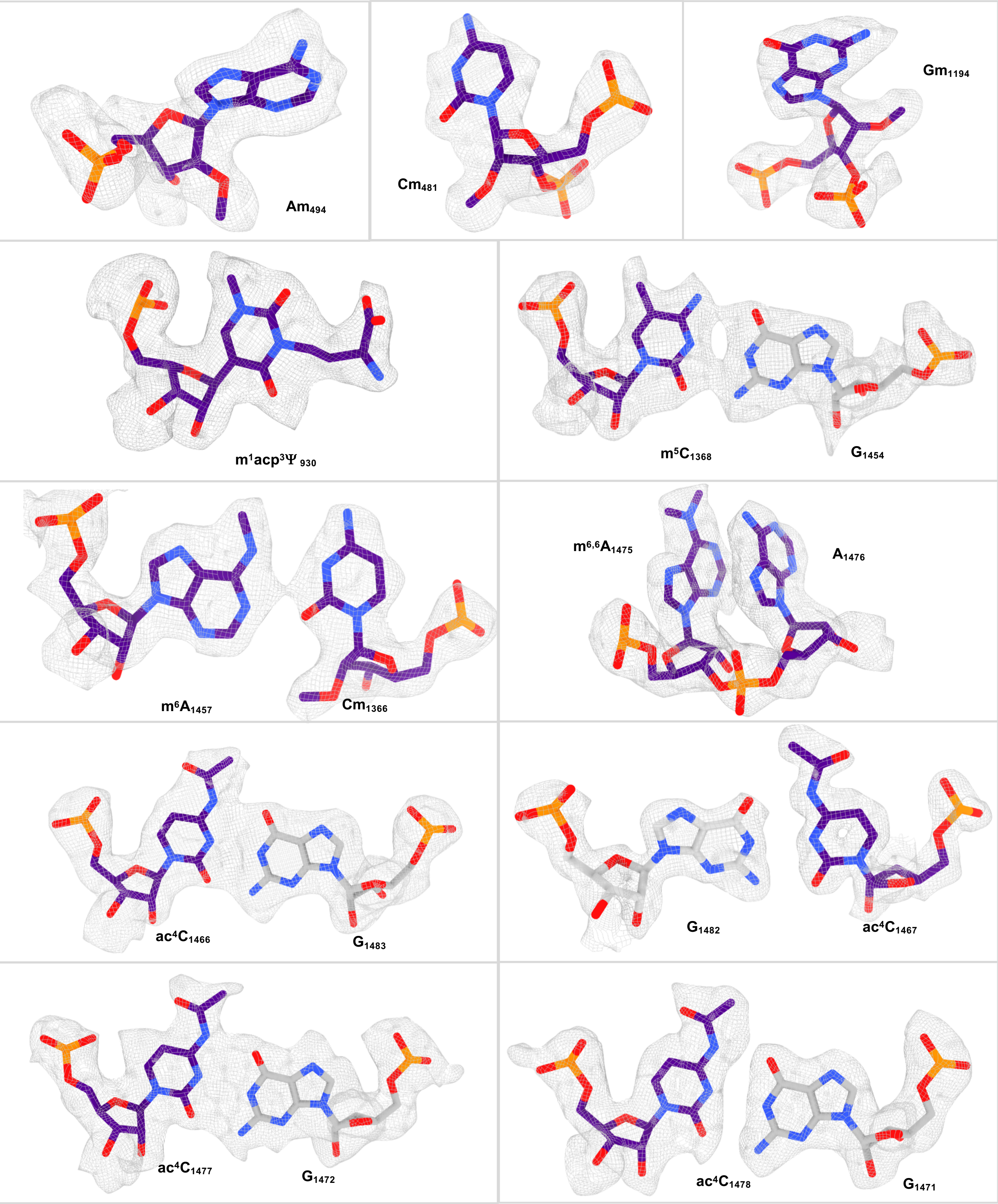
$Am_{494}$
$Cm_{481}$
$Gm_{1194}$
$m^{1}acp^{3}\Psi_{930}$
$m^{5}C_{1368}$
$G_{1454}$
$m^{6}A_{1457}$
$Cm_{1366}$
$m^{6,6}A_{1475}$
$A_{1476}$
$ac^{4}C_{1466}$
$G_{1483}$
$G_{1482}$
$ac^{4}C_{1467}$
$ac^{4}C_{1477}$
$G_{1472}$
$ac^{4}C_{1478}$
$G_{1471}$

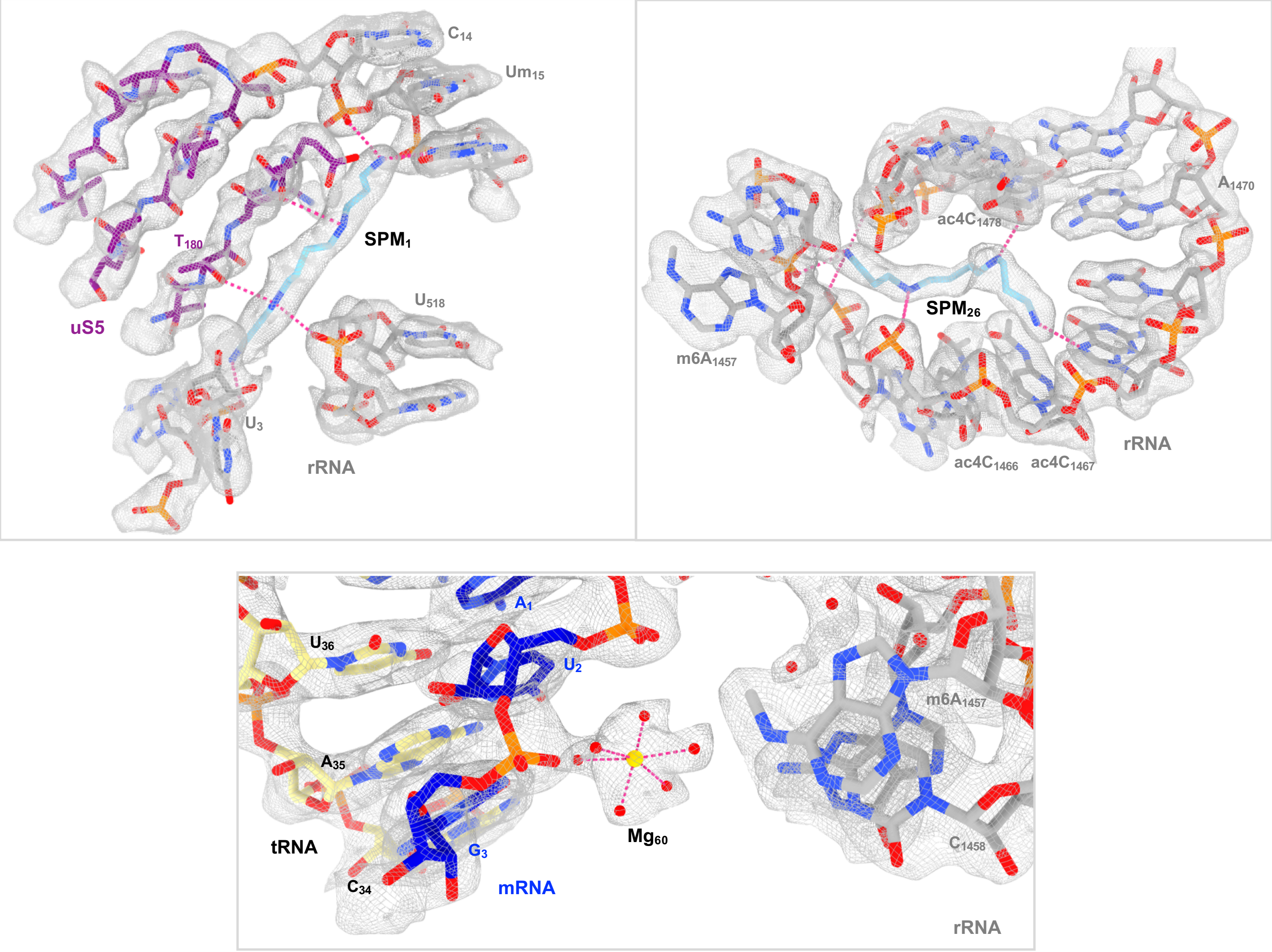

**Supplementary Figure 9: Examples showing the quality of the cryo-EM map for rRNA modified nucleotides, spermine residues, magnesium ions**

The 2.5 Å resolution cryo-EM map (DS2) is shown using the zone command in ChimeraX.

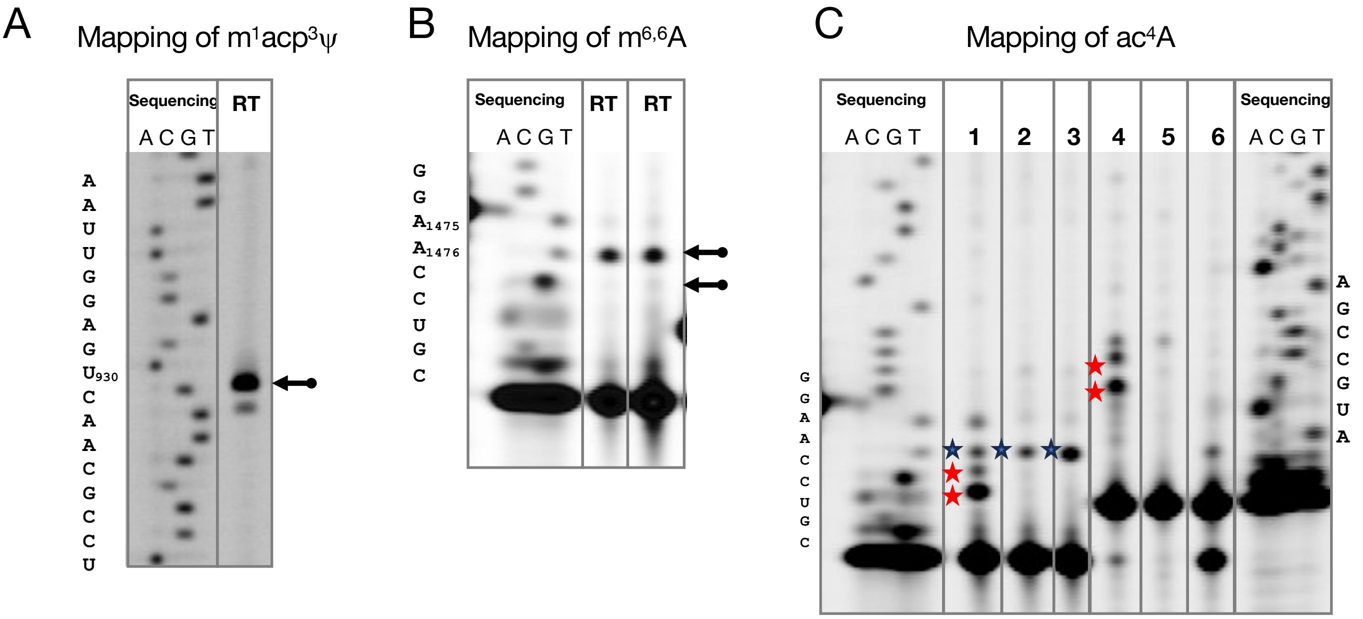

**Supplementary Fig. 10: Primer extension analysis of 16S rRNA**

(A) Mapping of the m¹acp³ψ modification expected at position 930. The arrow indicates the strong arrest of the reverse transcriptase one nucleotide before the expected modification. The weak arrest observed one nucleotide below could be due to the large steric hindrance of this modification. The sequence of the RT primer is 5'-GCAAGGTCGTTAGCCTGGCCG

(B) Mapping of the modifications $m^{6,6}$ expected at positions A1475 and A1476. The arrow indicates the strong arrest of the reverse transcriptase one nucleotide before the expected modification. The experiment shows the presence of a modification at position A1475. However, no convincing arrest signal showing the presence of an $m^{6,6}$ modification at position A1476 was observed. The sequence of the RT primer is 5'-GGAGGTGATCCAGCC.

(C) Primer extension analysis of 16S rRNA to map $ac^4C$ residues. Lanes 1 or 4, 16S rRNA treated with NaBH4 (100mM, 37°C, 1h as described in[28]), lanes 2 or 5 control 16S rRNA (37°C, 1h), lanes 3 or 6, intact 16S rRNA. The RT stops are indicated to the left of the bands with stars. Lanes 2 and 3 indicate the RT stop due to the presence of a dimethyl adenosine at position 1475. This stop occurs even in the presence of NaBH4 treatment. Lane 1 shows two additional arrests, as compared to lanes 2 and 3, indicative of ac4C in positions 1477 and 1478 (see also supplementary Figure 7). The sequence of the RT primer is 5'-GATCCAGCCGCAGGTTCC. Lane 4 indicates two additional arrests (red stars), as compared to lanes 5 and 6, indicative of ac4C in positions 1466 and 1467. The sequence of the RT primer is 5'-GGAGGTGATCCAGCC.

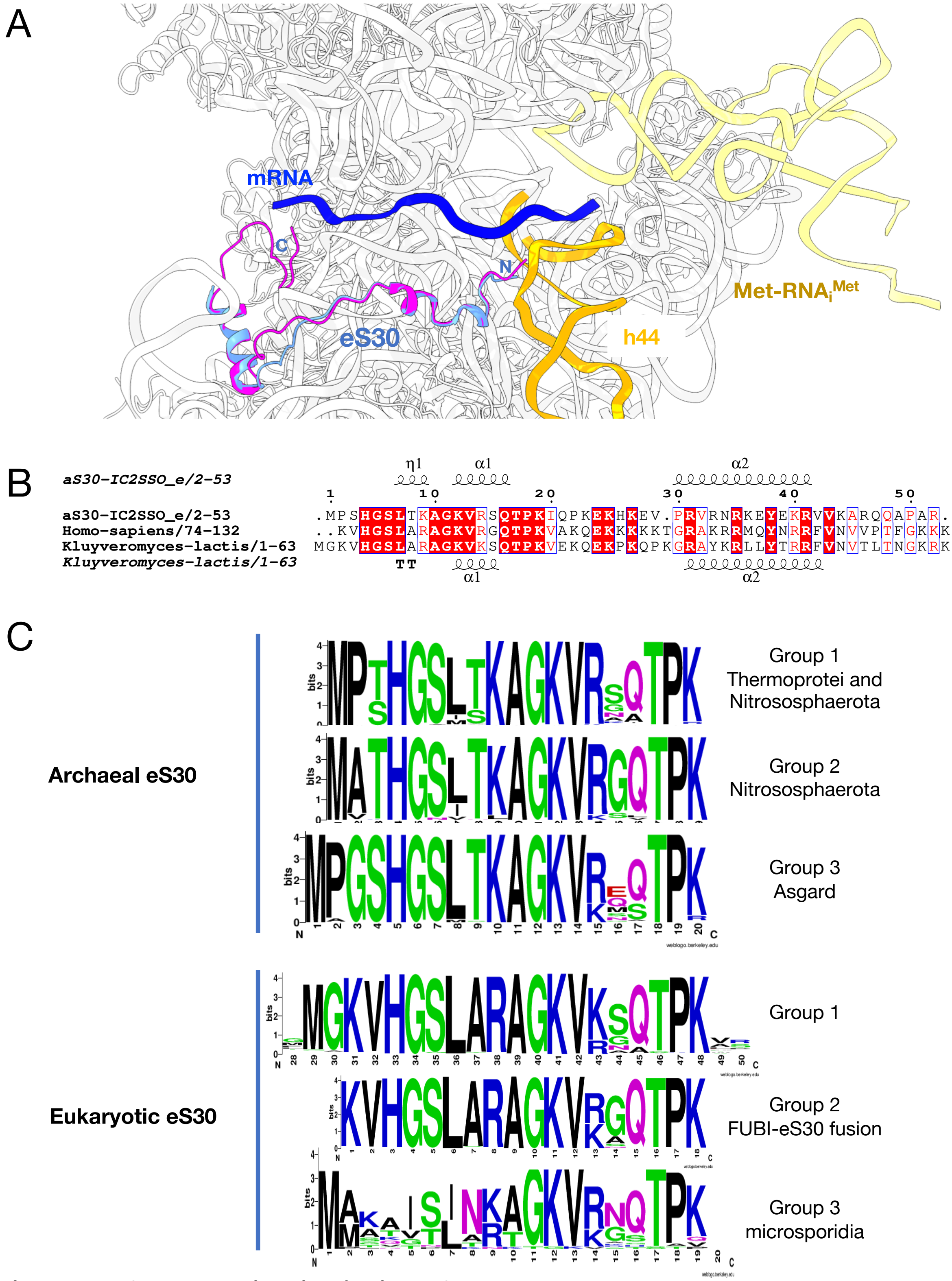

### Supplementary Figure 11: archaeal and eukaryotic eS30

A- Cartoon view of IC-DS2 with Ss-eS30 colored in cornflower blue. eS30 from superimposed human ribosome (8G5Z)[29] is colored in magenta. The view shows a remarkable conservation between archaeal and eukaryotic eS30 structures. Eukaryotic versions of eS30 have a longer C-terminal tail. Moreover, variability is observed at the N-terminal ends of the eukaryotic and archaeal proteins as shown in view B.

B-Archaeal and eukaryotic eS30 protein sequences were retrieved from Uniprot and aligned usingClustal Omega. The multi-sequence alignment was manually corrected using Jalview editor[30]. 703 archaeal sequences were aligned. The sequences corresponding to the first 23 amino acids were extracted. Three main groups were defined to calculate logos (https://weblogo.berkeley.edu/logo.cgi). Group 1 contains 522 sequences coming from Thermoprotei and Nitrososphaerota. Group 2 contains 84 sequences from Nitrososphaerota (all having MATH as first amino acids) with some Bathy-, Brock- and one Heimdall-archaeota. Group 3 contains 97 sequences from Asgard only (one exception for an unidentified thermococci) . Notably, Heimdallarchaeota have MAGSH instead of MPGSH. 2098 eukaryotic eS30 sequences manually curated were aligned. 830 sequences were identified as ubiquitin-like fusion protein FUBI-eS30. In these cases, the N-terminal extremity of eS30 obtained after cleavage of the FUBI-eS30 fusion protein was proposed to be the lysine of the KVHGS[31,32,33]. The remaining 1243 sequences corresponding to N-terminal regions were extracted and used to calculate a logo. The vast majority of sequences starts at position 29 of the multiple alignement. 25 sequences of microsporidia were considered separately. Collectively, this analysis shows that archaeal versions of eS30 have sequence specificities at the N-terminus as compared to eukaryotes. The archaeal version of eS30 that is closest to eukaryotes is that from Heimdallarchaeota.

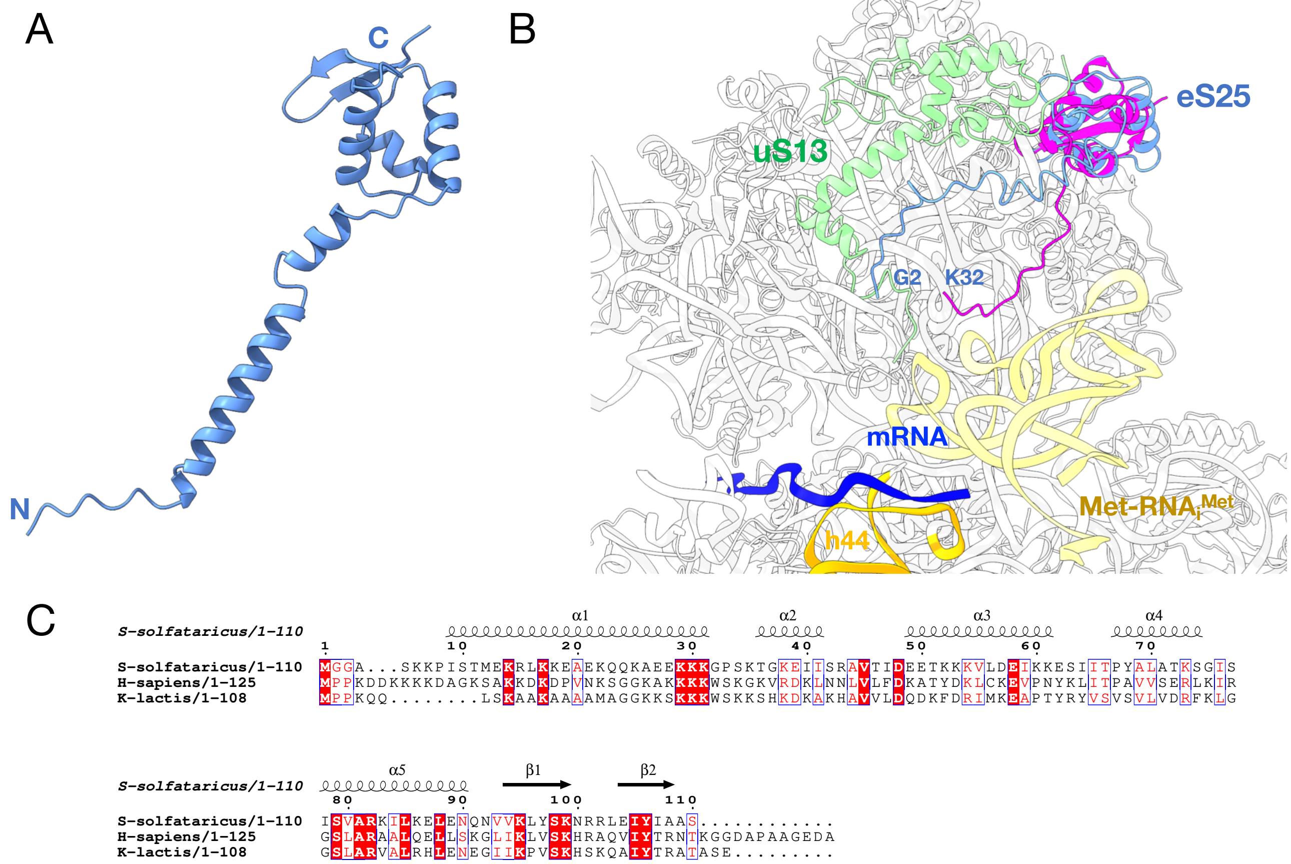

**Supplementary Figure 12**: **archaeal and eukaryotic eS25**

A. Ss-eS25 Alphafold2 model. The same prediction is available for eukaryotic and archaeal eS25.
B. Cartoon representation of IC-DS2 with Ss-eS25 colored in cornflower blue. Region 12 to 35 was only tentatively placed in a poorly defined part of the cryo-EM map. Human eS25 (PDB 8G5Z) is colored in magenta. Residues 1 to 32 of human eS25 are not visible.
C. Structural alignment of *S. solfataricus* eS25 with the human and *K. lactis* proteins. Notably, sequence alignment of 250 archaeal versions of eS25 show that about 70% of the sequences (most TACK except some Acidilobales, Desulfurococcales and Fervidicoccales) have an MGG sequence at the N-terminus.

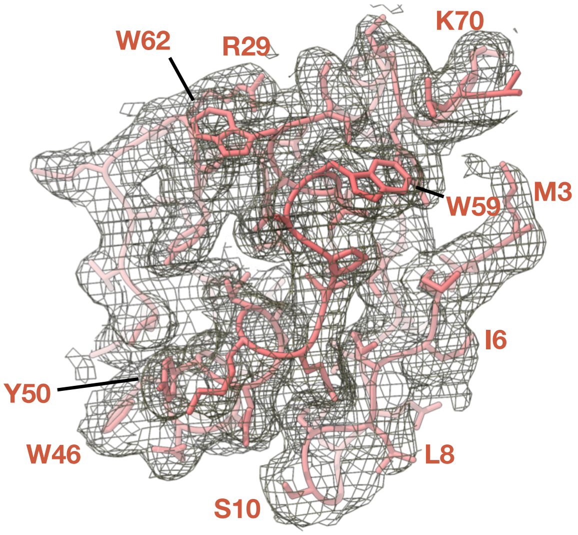

```
1       10        20        30        40        50        60        70
MEMLKIKLSSGREVEINDDTIAVLNEYVRTQMTLEELSKRLGLSGWEEAYELIKQVPAWVMWSPLPIYKKLA
```

**Supplementary Figure 13: aS33**

The view shows the quality of aS33 fitting into the cryo-EM map (multibody refined map EMDB-50445).

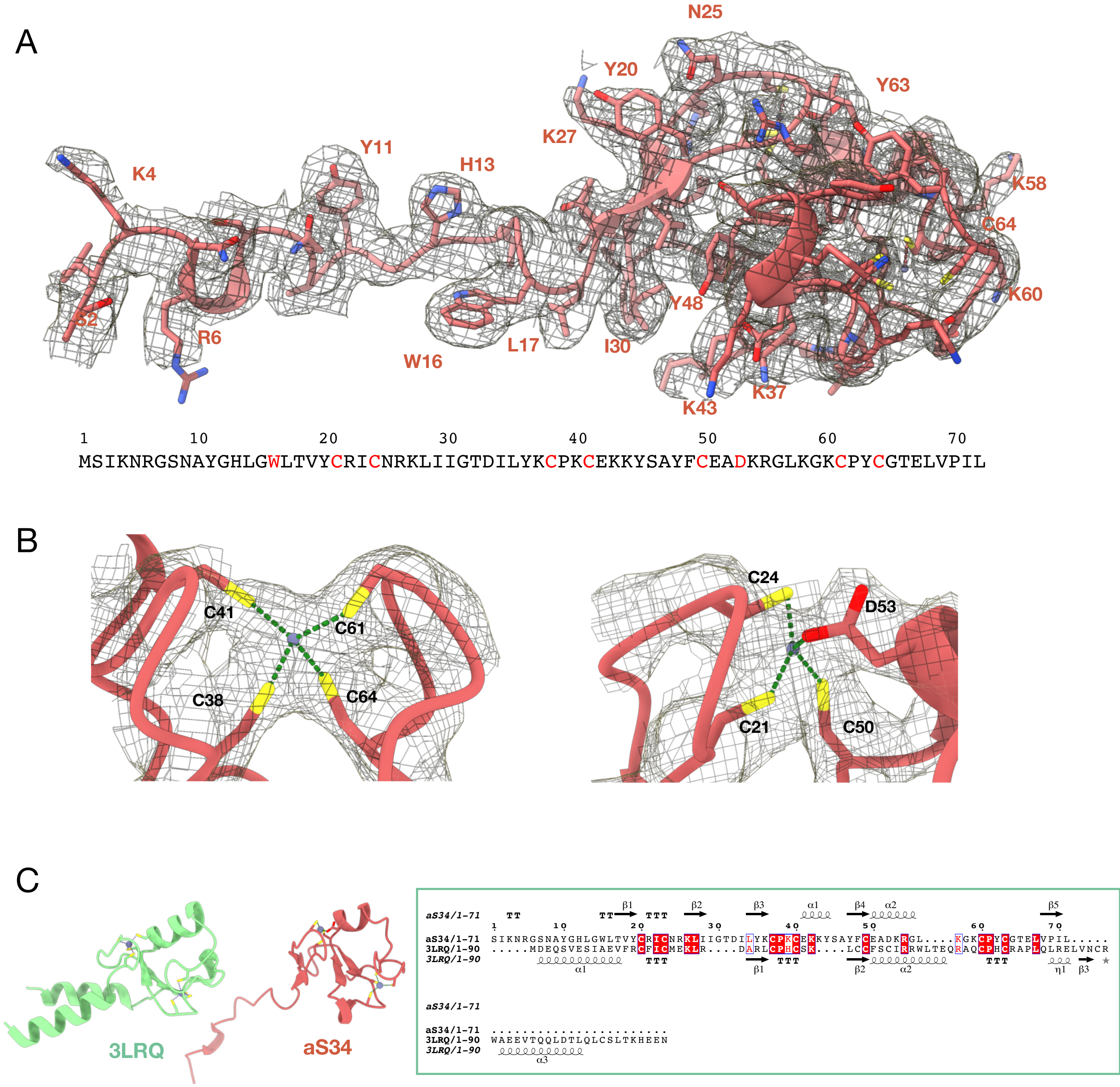

**Supplementary Figure 14: aS34**

A- The view shows the quality of aS34 fitting into the cryo-EM map (Local scale map EMDB 504445).
B-Close up views of the two zinc binding knuckles.
C- Superimposition of aS34 with human ubiquitin-protein ligase E3 RING domain (PDB 3LRQ). 3LRQ is in light green, aS34 in red. Rmsd 3LRQ to aS34, 2.9 Å for 217 atoms. Structural alignments were performed with Espript[14].

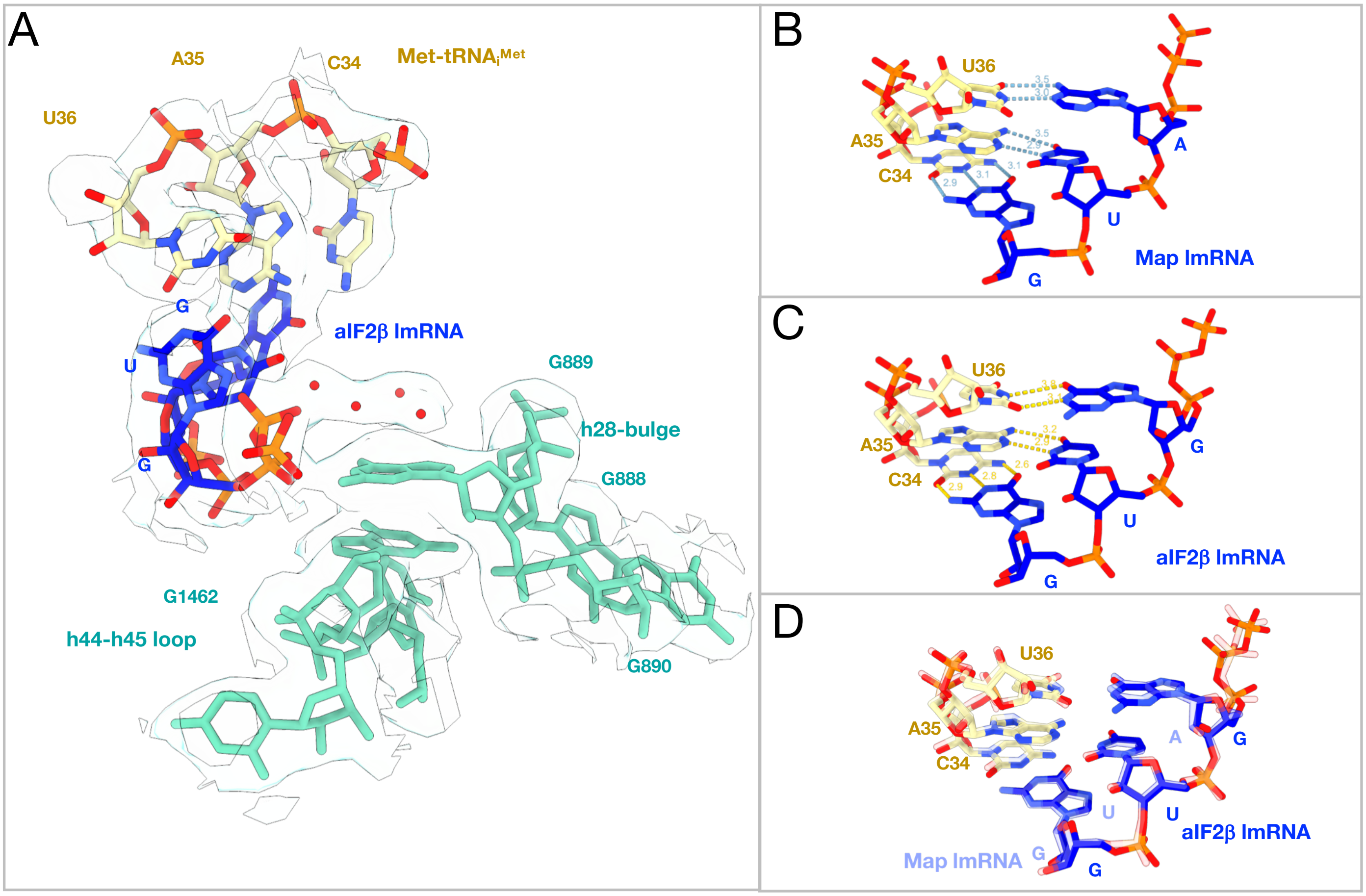

**Supplementary Figure 15: P site interaction in DS3 and comparison with DS2**

A. Close-up of the codon:anticodon interaction in IC-DS3. The cryo-EM map is represented to highlight the network of water molecules located above G889.
B. Close-up showing the AUG:CAU codon:anticodon interaction in IC-DS2.
C. Close-up showing the GUG:CAU codon:anticodon interaction in IC-DS3.
D. Superimposition of views B and C with view B shown in transparency. The view shows slight adjustment of the codon:anticodon interaction depending on the sequence of the start codon.

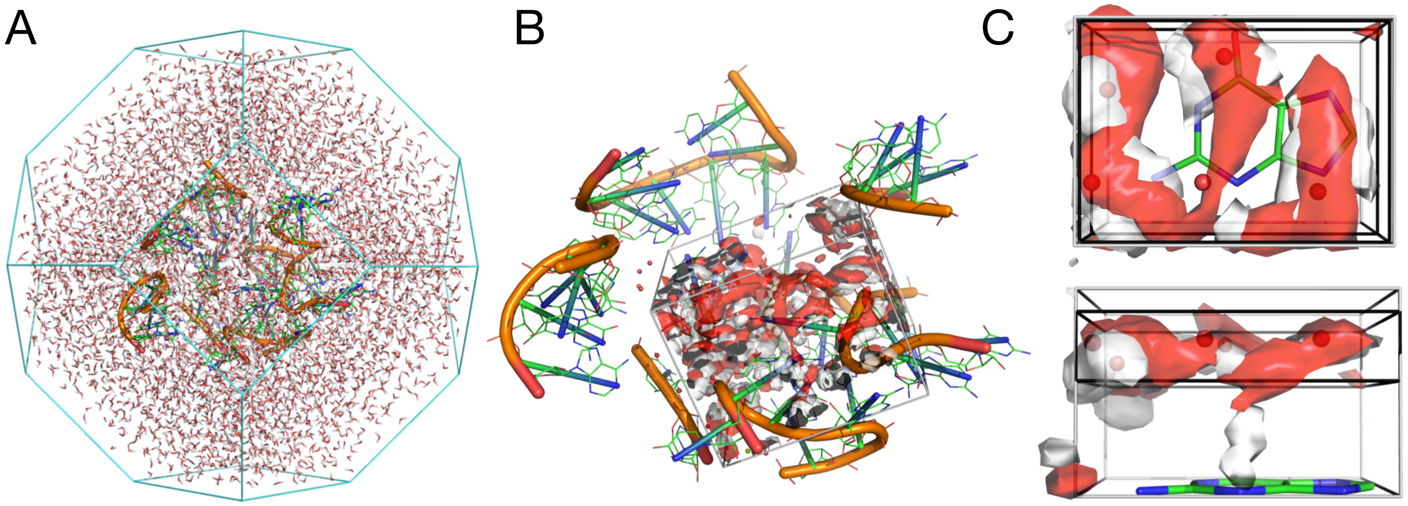

**Supplementary Figure 16: Explicit water molecular dynamics simulation**

A. Simulated system. The ribosomal molecular system was truncated at 12 Å around G889 nucleobase, immersed in a truncated octahedron box extending 10 Å away from the solute and filled with water molecules.
B. The view shows the simulated system along with water oxygen and hydrogen densities obtained by GIST.
C. Zoomed view on the region above G889 nucleobase. The view shows that the GIST predicted water oxygen (red) and hydrogen (white) densities fit with the experimentally placed water molecules (red spheres).

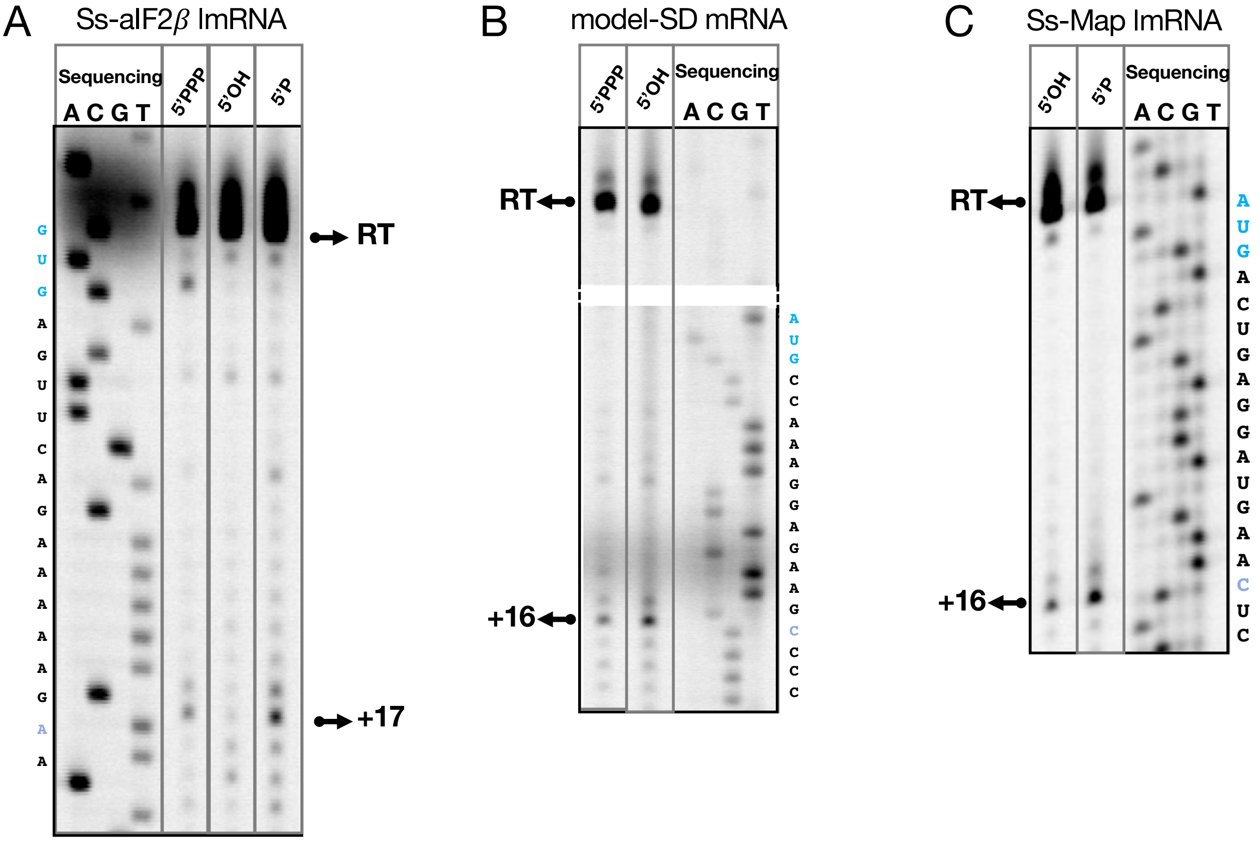

**Supplementary Figure 17: Toeprinting analysis of 30S complexes assembled on mRNAs with a 5'PPP, 5'OH or 5'P end**

A. Typical experiment using Ss-aIF2β lmRNA. Position of the initiation codon is indicated in blue. Position of the toeprinting signal observed at +17 is shown in light blue. The sequence of the mRNA complementary to the DNA sequence is shown on the left of the gel. Toeprinting signal obtained for a 30S:mRNA:Met-$\text{tRNA}_i^{Met}$:aIF2 complex where mRNA is 5'PPP, 5'OH, or 5'P, as indicated.

B. Same as panel A but using Model-SD mRNA.

C. Same as panel A but using Ss-Map lmRNA. This lmRNA was produced by *in vitro* transcription using a Hammerhead ribozyme construct (see Methods). It therefore has a 5'-OH end. Lane 1 is with a 5'OH end and lane 2 is with a 5'P end added with polynucleotide kinase.

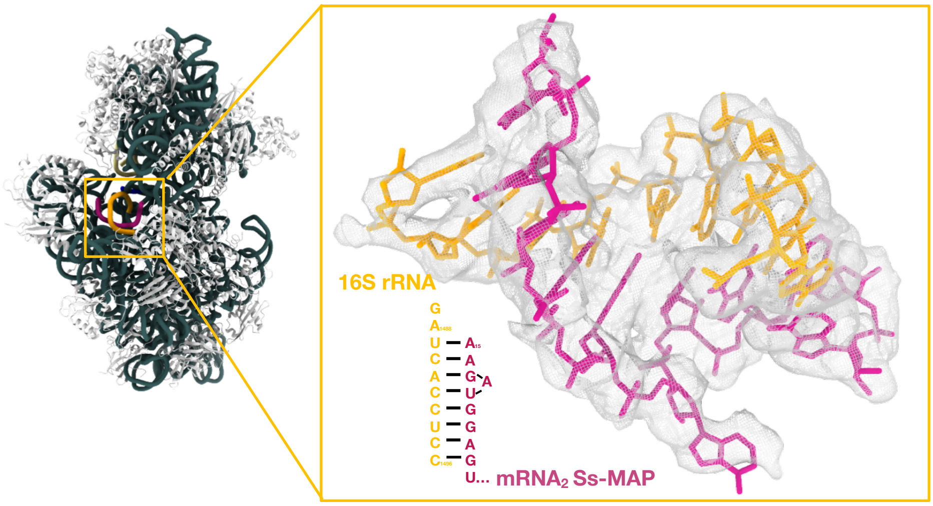

**Supplementary Figure 18: mRNA exit channel in IC-DS2**

The close up view shows the interaction of a second mRNA molecule (mRNA$_2$ Ss-MAP) with the anti SD sequence of the 16S rRNA.

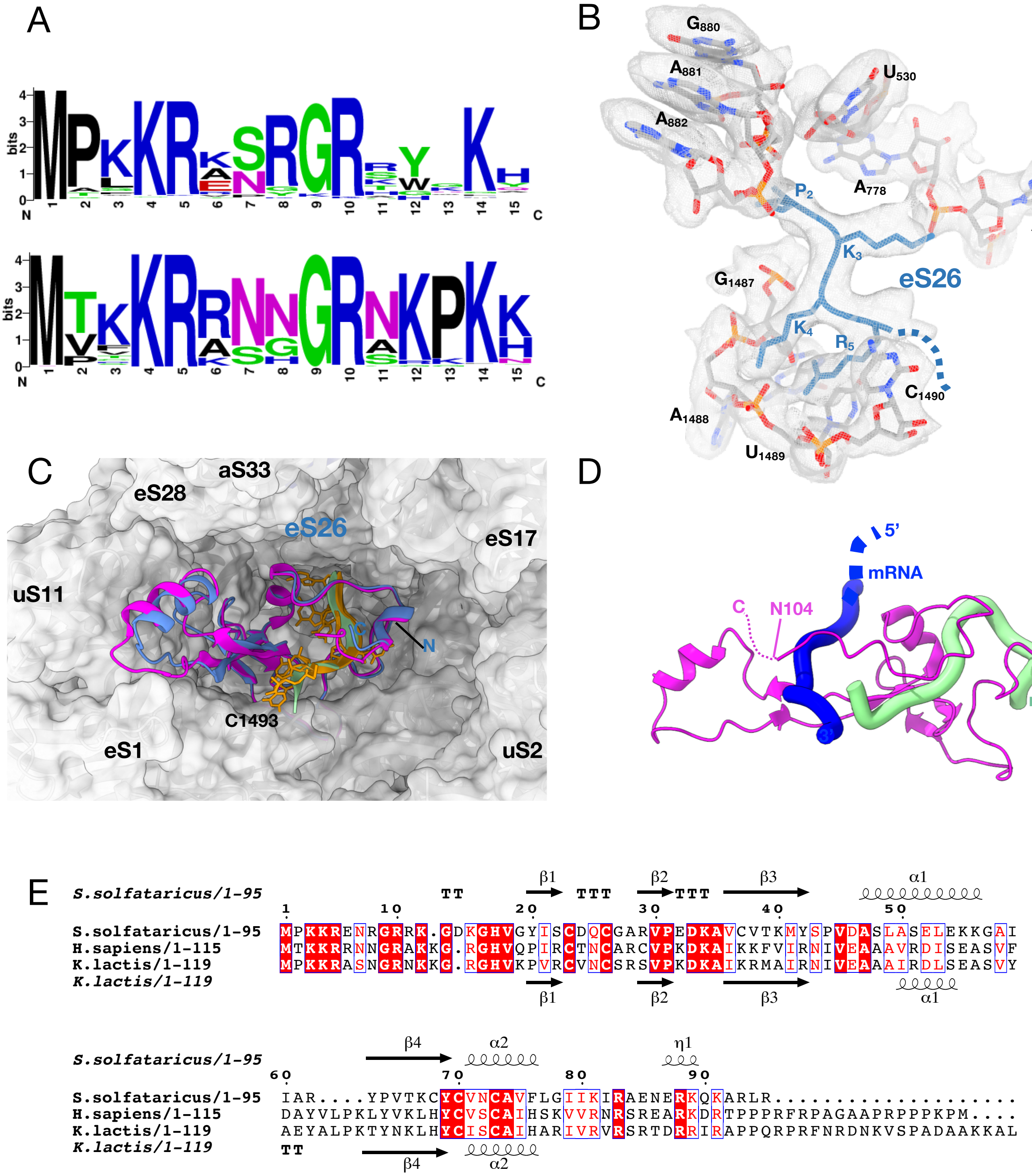

**Supplementary Figure 19: archaeal and eukaryotic eS26**

A-Archaeal and eukaryotic eS26 protein sequences were retrieved from Uniprot and aligned using Clustal Omega. The alignment was refined manually. 270 archaeal sequences were aligned. Average length of the protein is 95 amino acids. The sequence logo (Top) of the N-terminal residues shows a preference for the MPKKR sequence. The N-terminal peptide is tightly bound to the ribosome, as shown in view B. 1978 eukaryotic eS26 were aligned. 645 sequences containing unusual N-terminal regions were removed from the alignement. The calculated logo is shown at the bottom of the panel. Average length of the protein is 115 amino acids. As compared to archaea, eukaryotic eS26 have longer C-terminal tail containing low complexity regions rich in proline residues. The C-tail (residues 97 to 104) of *S. cerevisiae* eS26 contact the mRNA[27].

B-Binding of the Ss-eS26 N-terminal tail to 16s rRNA as observed in IC-DS3. The rRNA 3' end is shown in grey.

C- Structure of IC-DS3 with Ss-eS26 colored in blue. Human eS30 from 8G5Z is colored in magenta. Ss-16S rRNA extremity is in orange and that of human 18S rRNA is in light green. The view shows a remarkable conservation between archaeal and eukaryotic eS26 structures as well as in the binding to the rRNA

D-Binding of *S. cerevisiae* eS26 to rRNA and mRNA (PDB 6FYX)[27]. The view shows that the C-terminal tail of eS26 contacts the mRNA.

E- Structural alignment of *S. solfataricus* eS26 with the human and *K. lactis* proteins.

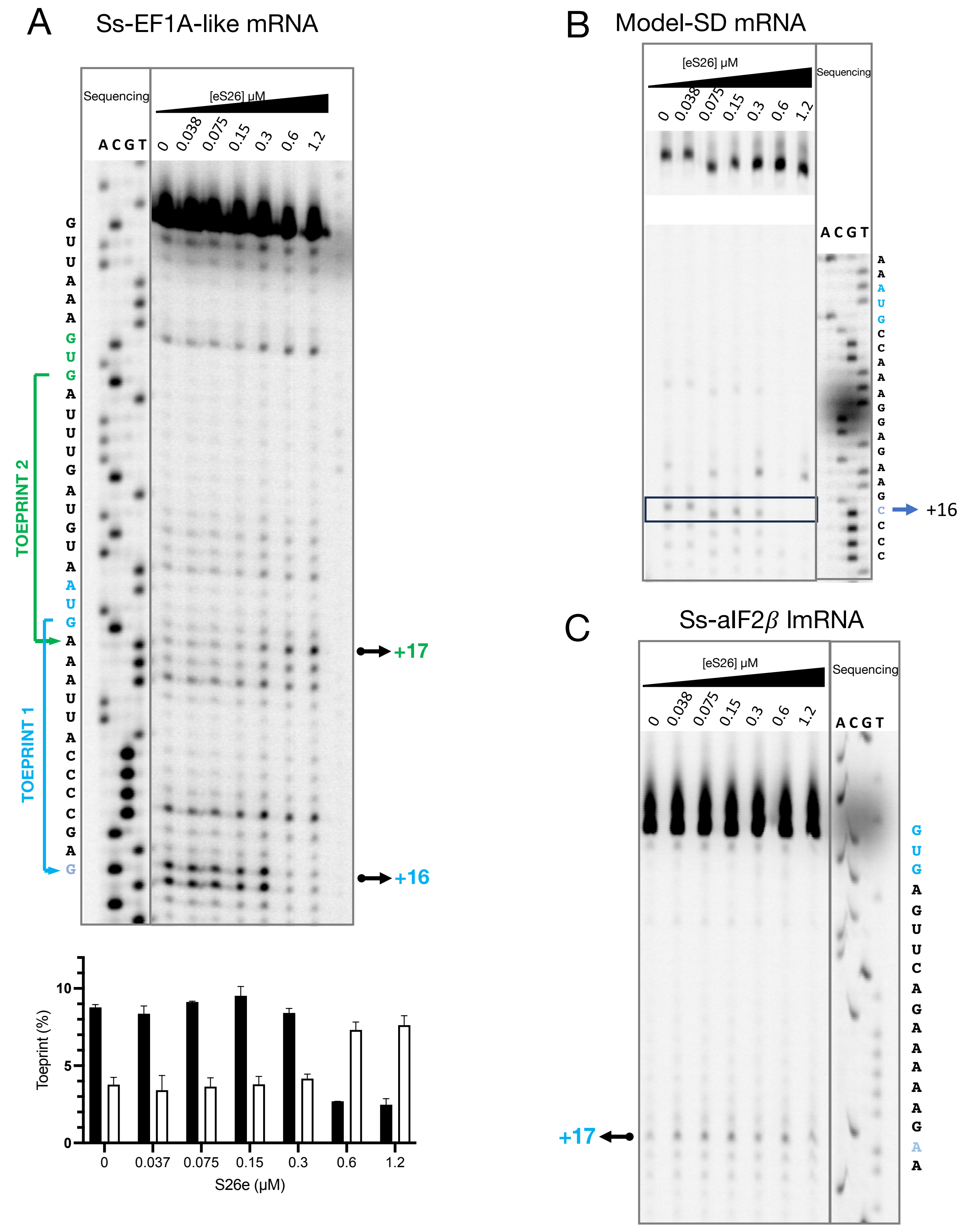

**Supplementary Figure 20: high concentrations of eS26 destabilize the initiation complexes formed on leadered mRNAs**

A. Influence of Ss-eS26 concentration on toeprinting signal intensity obtained with a 30S:mRNA:aIF2:Met-tRNA$_i^{Met}$ complex. The mRNA is Ss-EF1A-like. The two toeprinting positions discussed in the text are indicated. At high Ss-eS26 concentrations, the signal corresponding to toeprint 2 increases. Below the gel image is the quantification of the main band corresponding to toeprint 1 or 2.
B. Same as A but for Model-SD mRNA.
C. Same as A but for Ss-aIF2β lmRNA.
